# Mode of T cell priming durably shapes the TCR repertoire, effector function and α4β1 integrin expression of human virus-specific CD4^+^ T cells

**DOI:** 10.64898/2026.08.18.745226

**Authors:** Elie Antoun, Guihai Liu, Deshni Jayathilaka, Xuan Yao, Timothy Rostron, Craig Waugh, Kevin Clark, Paul Sopp, Jeremy W. Fry, Tianyi Xia, Alexander J. Mentzer, Julian C. Knight, Yanchun Peng, Tao Dong

## Abstract

The generation of an effective T cell response against an antigen depends on the recognition of the antigen by the T cell receptor (TCR), followed by T cell priming, initiating coordinated biophysical, biochemical and proliferative changes that drive differentiation into effector and memory clones. The immunological environments in which priming occurs, such as natural infection or vaccination, influences the quality and persistence of memory T cells, but the long-term impacts remain incompletely understood. Here, we investigate how the mode of priming shapes durable antigen-specific CD4^+^ T cell memory, utilising two cohorts 3-4 years after initial antigen encounter: individuals recovered from SARS-CoV-2 infection and infection-naive individuals who received a SARS-CoV-2 vaccination. Using ex vivo single-cell RNA sequencing, paired TCR sequencing and *in vitro* functional analyses, we characterise the transcriptional, clonal and functional profile of Spike-specific CD4^+^ T cells. Across both cohorts, CD4^+^ T cell responses against spike epitopes S_166-180_, S_751-765_ and S_866-880,_ were immunodominant, with shared public TCR clonotypes indicating conserved antigen-recognition regardless of mode of priming. Despite this shared specificity, infection-primed individuals exhibited greater TCR repertoire diversity and lower CDR3αβ sequence convergence. Transcriptionally, infection-primed cells exhibited a more cytotoxic and effector phenotype, while vaccine-primed cells preferentially adopted T follicular helper (Tfh)- and Th1-associated phenotypes. Infection-primed individuals also displayed enrichment of cell adhesion and integrin signalling pathways, with a greater proportion of spike-specific CD4^+^ T cells expressing α4β1 integrin subunits, consistent with enhanced migratory and effector potential. Collectively, our findings demonstrate that the mode of antigen priming may influence the long-term CD4^+^ T cell memory states, influencing TCR repertoire diversity, functional differentiation and tissue-homing potential, years after the initial immune response.

## Introduction

The generation of an effective adaptive immune response depends on the successful priming of naïve T cells by primary recognition of peptide-major histocompatibility (pMHC) complexes on antigen-presenting cells, which consists of range of biophysical, biochemical and proliferative changes leading to expanded and differentiated effector clones^1^. During T cell priming, T cells integrate signals derived from pMHC recognition by the T cell receptor (TCR), co-stimulation by cell surface co-receptors and signalling from inflammatory cytokines such as IL12^2^ and interferons^3^ to initiate programmes of proliferation, differentiation, and memory formation. These priming events not only determine the magnitude of the primary immune response but also influence the long-term characteristics of the resulting memory T cell population. A growing body of evidence suggests that the quality of T cell priming can shape the overall T cell differentiation trajectories and memory cell fates, with lasting consequences for protective immunity^1,4–7^.

Natural infection and vaccination represent two distinct modes of T cell priming. Infection typically exposes the immune system to the infectious pathogen at the site of infection (e.g. respiratory tract for Influenza A/SARS-CoV-2 or the skin and mucous membranes for herpes simplex virus), and is accompanied by a complex innate immune activation and prolonged antigen exposure^8^. In contrast, vaccination delivers antigen in a more controlled context, often subcutaneously or intramuscularly, with restricted tissue distribution, where antigen-presenting cells subsequently migrate to the lymph nodes for T cell priming^9^. Vaccine administration results in a fundamentally different inflammatory environment. These differences are known to influence the overall magnitude and phenotype of primary T cell responses. However, whether the initial mode of priming leaves a durable long-term imprint on antigen-specific CD4^+^ T cell memory remains incompletely understood.

Borcherding et al.^10^ have shown that CD4^+^ T cells in both the draining lymph nodes and the circulation exhibit distinct transcriptional phenotypes following either SARS-CoV-2 vaccination or infection. Spike-specific CD4^+^ T cells showed a predominant BCL2^+^CD4^+^ T_CM_ phenotype, while those from vaccinated individuals showed a greater expression of pro-survival and antiapoptotic gene signatures, suggestive of improved long-term survival. However, spike specificity in this study was inferred using Trex, rather than experimentally determined. Furthermore, the samples were obtained from individuals up to 6 months after vaccination, and so whether there is improved long-term survival of these cells in vaccinated individuals is unclear.

Given the importance of the role of T cell priming in the phenotype and function of T cells, the long-term consequences of the mode of priming is likely reflected in both the TCR repertoire and transcriptional phenotype of the antigen-specific memory T cell compartment. Although studies in animal models have demonstrated that inflammatory signals encountered during priming can alter both TCR selection and memory differentiation, evidence for persistent priming-dependent effects on human antigen-specific CD4^+^ T cells remains limited.

The recent SARS-CoV-2 pandemic provided a unique opportunity to address this question. For the first time, large populations of individuals acquired immunity to a previously novel pathogen through either natural infection or vaccination in infection-naïve individuals, allowing direct comparison of memory T cells generated against the same antigen but under fundamentally different priming conditions, 3-4 years after initial priming. Here, we used ex vivo single-cell transcriptomic, paired TCR repertoire, and *in vitro* functional analyses to examine CD4^+^ T cells specific for three immunodominant SARS-CoV-2 spike epitopes in individuals whose responses were initially primed either by infection or vaccination. By analysing antigen-specific memory CD4^+^ T cells several years after the initial antigen encounter, we sought to determine whether the mode of priming leaves a persistent influence on the TCR repertoire and phenotype of antigen-specific CD4+ T cells. Our findings demonstrate that infection- and vaccine-mediated priming generate distinct long-term memory populations characterised by differences in TCR repertoire organisation, cytotoxic potential, and integrin-associated migratory programmes, revealing a lasting influence of the priming environment on human CD4^+^ T cell memory.

## Results

### 1. Immunodominant CD4^+^ T cell responses to SARS-CoV-2 spike protein in the vaccine-primed and infection-primed cohorts

We and others have previously identified and reported on three dominant SARS-CoV-2 spike protein (S) CD4^+^ T cell epitopes: S_166-180_ (CTFEYVSQPFLMDLE)^11–13^, S_751-765_ (NLLLQYGSFCTQLNR)^11,14^ and S_866-880_ (TDEMIAQYTSALLAG)^11,15,16^, restricted by HLA-DPB1*04:01 for S_166-180_ and by HLA-DRB1*15:01 for S_751-765_- and S_866-880_-specific T cells. We have previously investigated these T cells in our previously described convalescent cohort of individuals primed by natural infection^17^.

To determine whether all three of these immunodominant responses were similarly induced by vaccination, we recruited 68 healthy volunteers from early 2021, following the rollout of SARS-CoV-2 vaccination in the UK (Supplemental Figure 1a). Participants received either the ChAdOx1 vaccine or an mRNA vaccine, and vaccine-induced Spike-specific T cell responses were longitudinally monitored following multiple vaccine doses. All participants underwent HLA-DR and HLA-DQ typing (Supplementary Table 1). 20 individuals were HLA-DRB1*15 positive, while HLA-DP alleles were not determined. *Ex vivo* IFN-γ ELISpot assays were performed to assess T cell responses against epitope S_166-180_, S_751-765_ and S_866-880._ Amongst the 68 donors, 35 (54.4%) responded to S_166-180_, 17 (25%) to S_751-765,_ and 13 (19.1%) to S_866-880_ (Fig 1a). These response frequencies were broadly comparable to those observed in our convalescent cohort (52.5%, 32.4% and 27.8%, respectively, Fig. 1a), further confirming the immunodominance of these three Spike-derived CD4^+^ T cell epitopes and demonstrating that robust T cell responses against them can be effectively primed by SARS-CoV-2 vaccination.

**Fig. 1.**
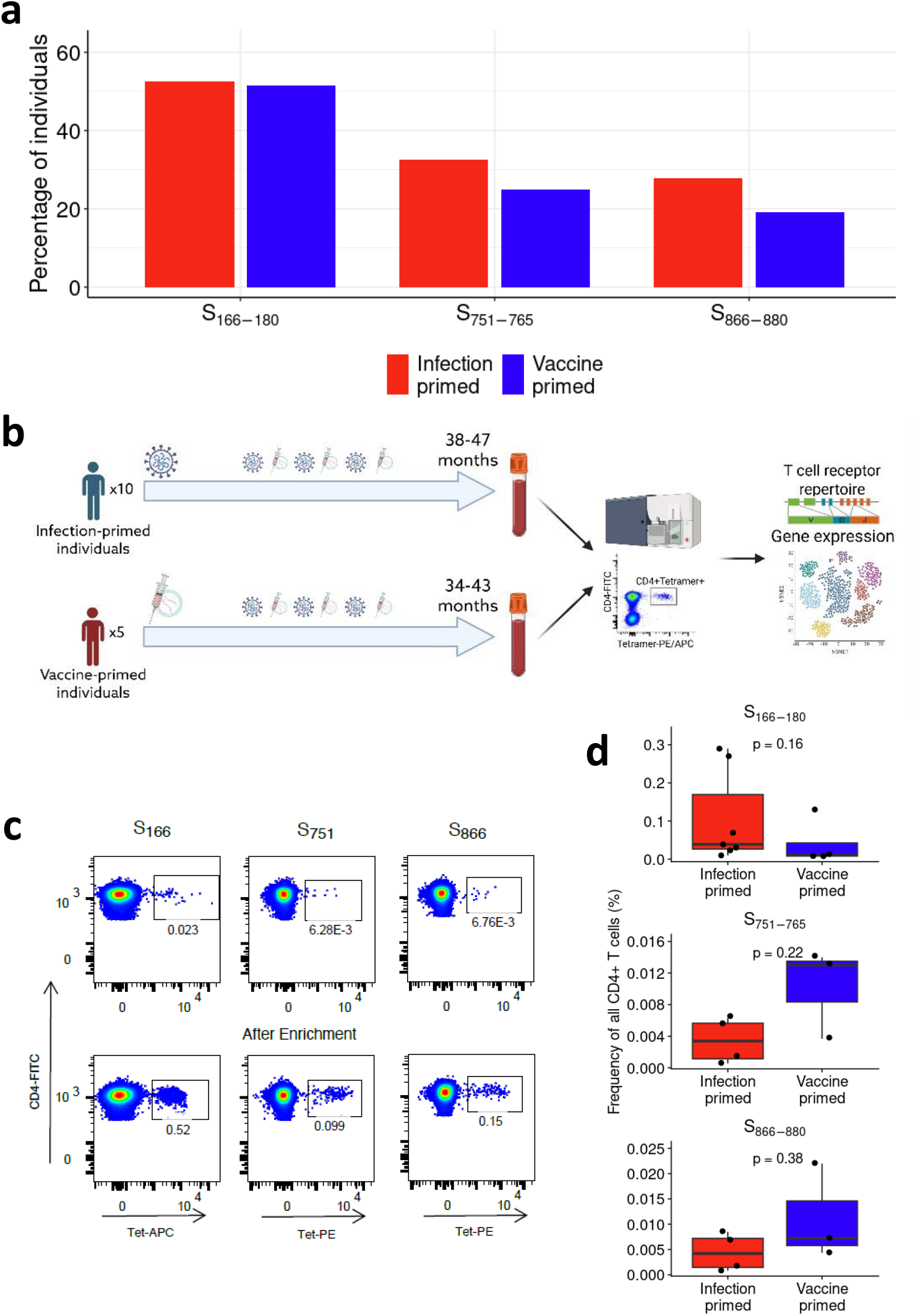
Overall study design and TCR repertoire characteristics. (a) Percentage of individuals in the infection-primed (red) and vaccine-primed (blue) cohort that elicited a detected IFN-g ELISpot response in response to S_166-180_, S_751-765_ and S_866-880_ peptides. Total individuals tested: n=68 vaccine-primed, n=40 S_166-180_ infection-primed, n=37 S_751-765_ infection-primed and n=36 S_866-880_ infection-primed (b) Overall study workflow, created in Biorender. (c) Representative gating of tetramer+ cell populations ex vivo (top panels) and after magnetic enrichment for tetramer+ cells (bottom panels). (d) Frequency of tetramer^+^ cells of all CD4^+^ T cells per individual. Wilcoxon signed-rank test used for pairwise comparisons in d.

To investigate long-term impact of natural infection and vaccination on antigen-specific CD4^+^ memory T cells, we recruited 14 individuals for detailed analysis: 9 individuals from convalescent cohort who had recovered from COVID-19 following primary infection in 2020 and were subsequently exposed through multiple dose of vaccinations and/or reinfection, and 5 individuals from the vaccination cohort who had been initially primed by vaccination, and subsequently had breakthrough infections (Supplementary Table 2). Participants were sampled 3-4 years after the initial priming event, defined as either primary infection or receipt of 1^st^ vaccine dose. T cell responses to these three immunodominant CD4 spike epitopes were characterised and an overview of the study design is shown in Fig. 1b.

Given the low ex vivo frequency of antigen-specific T cells, peptide-MHC II tetramer enrichment was performed prior to cell sorting, as previously described^18,19^. This approach increased the frequency of tetramer+ T cells by 20-30-fold, enabling downstream single-cell RNA Sequencing (scRNAseq) (Fig 1c, Supplemental Figure 1b-c). Of the individuals we recruited, 11 individuals (4 vaccine-primed and 7 infection-primed) elicited an S_166-180_-specific CD4^+^ response and stained positive with the HLA-DPB1*04:01 tetramer, while 8 individuals (3 vaccine-primed, 5 infection-primed) elicited an S_751-765_-specific and S_866-880_-specific CD4^+^ response and stained positive with their respective HLA-DRB1*15:01 tetramer (Supplementary Table 2). The frequencies of tetramer^+^ CD4^+^ T cells did not differ significantly between infection-primed and vaccine-primed individuals for any of the three epitopes (Fig. 1d, S_166-180_-specific p=0.16, S_751-765_-specific p=0.22, S_866-880_-specific p=0.38).

### 2. Public TCR clonotypes shared between vaccine-primed and infection-primed T cells

Tetramer^+^ CD4^+^ T cells were sorted, and *ex vivo* SmartSeq2 was performed to analyse the gene expression and paired TCR sequences from these spike-specific CD4^+^ T cells from the 15 individuals described above. In total, our TCR repertoire dataset comprised 1309 tetramer-sorted cells from 10 individuals primed by infection (863 S_166-180_-specific, 207 S_751-765_-specific and 239 S_866-880_-specific CD4^+^ T cells) and 760 tetramer-sorted cells from 5 individuals primed by vaccination (349 S_166-180_-specific, 224 S_751-765_-specific and 187 S_866-880_-specific CD4^+^ T cells).

Consistent with previous reports^13,20–22^, we observed a high diversity in V gene usage for both the alpha and beta chains (Fig. 2a-b). Overall, there was a broad V gene usage across all three epitopes for the TCRα (Fig. 2a) and TCRβ (Fig. 2b) chains. However, differences were evident based on the individuals’ mode of priming with particular V gene usages found at a higher frequency in on group of individuals (e.g. TRAV12-1 in a greater proportion of cells from vaccine-primed individuals, Fig. 2a middle panel), or in only one group of individuals (e.g. TRBV14 in infection-primed individuals, Fig. 2b right panel).

**Fig. 2.**
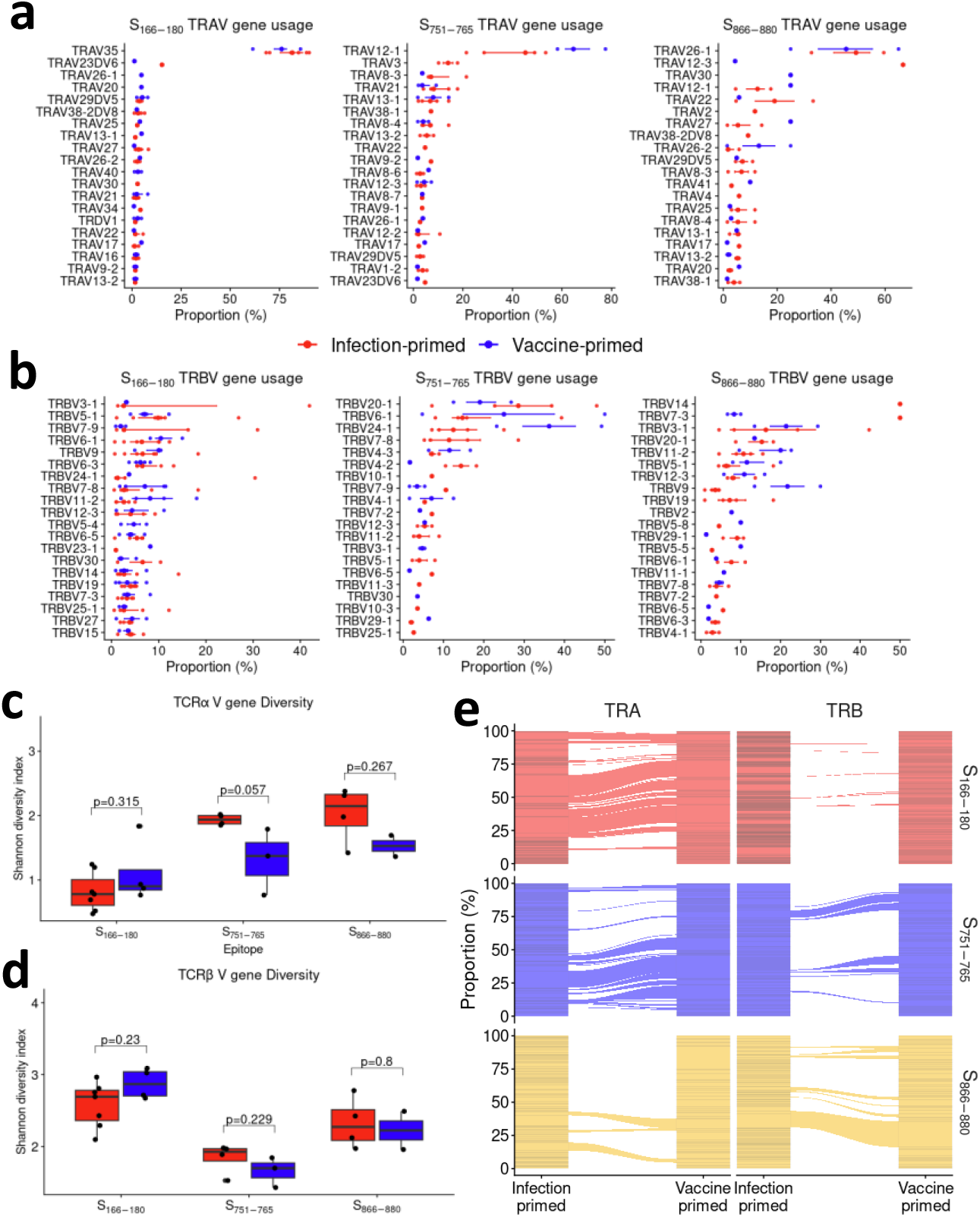
Global TCR repertoire analysis between infection- and vaccine-primed individuals. (a) Top 20 TRAV gene usage across epitopes, split by mode of priming, red=infection-primed, blue=vaccine-primed. (b) Top 20 TRBV gene usage across epitopes, split by mode of priming, red=infection-primed, blue=vaccine-primed. (c) Shannon diversity index comparing infection-primed (red) vs vaccine-primed (blue) individuals for TRAV gene usage, across the three specificities. (d) Shannon diversity index comparing infection-primed (red) vs vaccine-primed individuals for TRBV gene usage, across the three specificities. (e) Clonotype sharing between infection and vaccine primed individuals (CDR3+V gene). Wilcoxon signed-rank test used for pairwise comparisons in c and d.

Comparing the diversity of the V gene usage between the 3 epitopes. Although we see no statistically significant difference, comparing the TRAV and TRBV gene diversities between modes of priming, we can see trends for S_751-765_- and S_866-880_-specific cells showing reduced diversity of TRAV gene usage for vaccine-primed individuals compared to infection-primed individuals (Fig. 2c+d), while S_751-765_-specific cells also show reduced diversity in TRBV gene usages in vaccine-primed individuals. Moreover, S_166-180_-specific cells show a greater diversity of TRAV and TRBV gene usages for individuals primed by vaccination compared to infection-primed individuals (Fig. 2c+d). Furthermore, we identify individual chain sharing between vaccine-primed and infection-primed individuals, highlighting public shared clonotypes regardless of mode of priming (Fig. 2e), although there remains a significant level of TCR chains that are unique to infection-primed and vaccine-primed individuals, likely contributing to the repertoire differences.

### 3. S_751-765_-specific CD4^+^ T cells show different TCR repertoire properties, dependent on mode of priming

We next interrogated the properties of the TCR repertoire to determine whether mode of priming results in differences in the global repertoire. PCA analysis of the TRAV gene usage showed clear separation on PC1 corresponding to the HLA-restriction of the TCRs, with S_166-180_-HLA-DPB1*04:01-restricted T cells grouping separately to S_751-765_-specific and S_866-880_-specific T cells which are restricted by HLA-DRB1*15:01 (Fig. 3a). Furthermore, for HLA-DRB1*15:01-restricted T cells, cells from vaccine-primed individuals separate from cells from infection-primed individuals on PC2, with the greatest separation for S_751-765_-specific T cells.

**Fig. 3.**
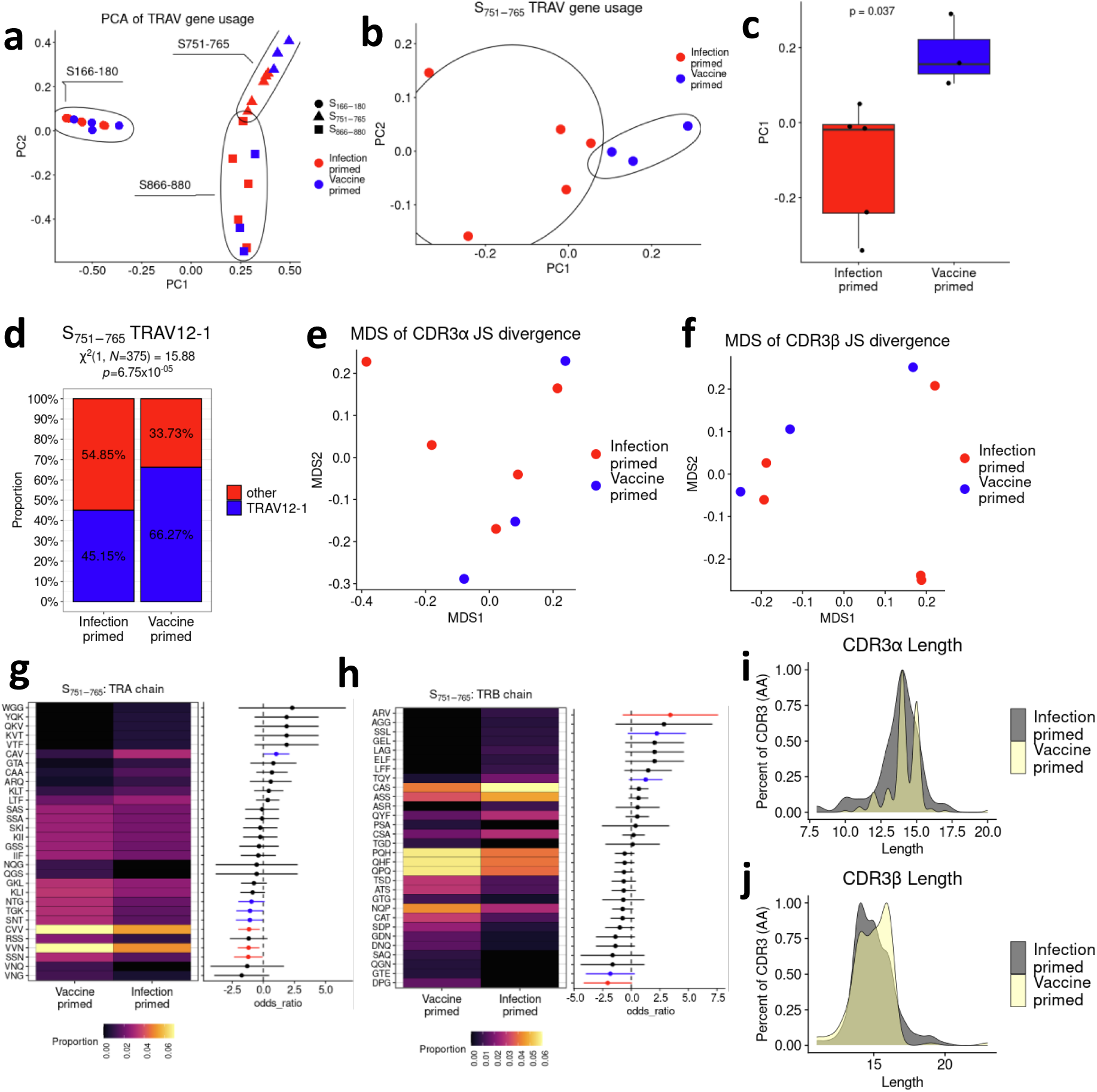
TCR repertoire comparison of S_751-765_-specific cells CD4+ T cells. (a) PCA of TRAV gene usage across all cells, coloured by mode of priming (red=infection-primed, blue=vaccine-primed), and circled by specificity. (b) PCA of TRAV gene usage across S_751-765_-specific cells, coloured by mode of priming (red=infection-primed, blue=vaccine-primed) (c) Box and whisker plot of PC1 values, grouped by mode of priming, showing median±IQR. P-value from a Wilcoxon signed rank test (d) Proportion of TRAV12-1+ cells (blue) or other TRAV usages (red) in cells from infection-primed and vaccine-primed individuals; χ^2^ test of independence to compare between groups. (e) MDS plot of Jensen-Shannon divergence of CDR3α sequences (f) MDS plot of Jensen-Shannon divergence of CDR3β sequences (g) Kmer heatmap of CDR3α sequences showing proportion of cells with specific 3mer sequences (left) and log2 odds ratio of Kmer enrichment (right). Fisher’s exact test used to identify significant enrichment with red lines=p<0.05 and blue lines=p<0.1. (h) Kmer heatmap of CDR3β sequences showing proportion of cells with specific 3mer sequences (left) and log2 odds ratio of Kmer enrichment (right). Fisher’s exact test used to identify significant enrichment with red lines=p<0.05 and blue lines=p<0.1. (i) Density plot of CDR3α length distributions between infection-primed (grey) and vaccine-primed (yellow) individuals. (j) Density plot of CDR3β length distributions between infection-primed (grey) and vaccine-primed (yellow) individuals.

We next further investigated the TCR repertoire of S_751-765_-specific T cells. PCA analysis of TRAV gene usage of S_751-765_-specific cells shows clear separation on PC1 of cells from vaccine-primed and cells from infection-primed individuals (Fig. 3b), with a greater PC1 for cells from vaccine-primed individuals (Fig. 3c, p=0.037). In addition, vaccine-primed samples formed a relatively tight cluster, whereas infection-primed samples were more broadly dispersed, consistent with greater heterogeneity in TRAV gene usage. Examining the feature loadings for PC1, TRAV12-1 shows the greatest positive contribution to PC1, and investigating the proportion of cells with TRAV12-1 gene usage, a greater proportion of cells from vaccine-primed individuals express TRAV12-1, compared to cells from infection-primed individuals (Fig. 3d, χ^2^ p=6.75×10^−5^). A similar association was seen for TRBV gene usage (Supplemental figure 2a), with a trend for higher PC2 in vaccine-primed individuals (Supplemental figure 2b), driven by a greater proportion of TRBV21-1+ cells in vaccine-primed individuals (Supplemental figure 2c).

We next investigated the CDR3 sequences of the S_751-765_-specific T cells. To systematically compare the TCRs on the basis of sequence similarity, we the calculated pairwise Jensen-Shannon divergence between CDR3 probability distributions. Multidimensional scaling (MDS) visualisation projected the TCRs in a 2D space where proximity reflects shared CDR3 usage. For both the CDR3α (Fig. 3e) and CDR3β (Fig. 3f), there are subtle, but consistent sharing patterns of the CDR3 usage, with greater sharing amongst cells from vaccine-primed individuals compared to infection-primed individuals. Similar patterns of JS divergence are seen for CDR3α sequences of S_866-880_-specific and S_166-180_-specific CD4+ T cells (Supplemental figure 2d+e) and CDR3β sequences (Supplemental figure 2f+g), albeit to a lower extent. Kmer analysis of the CDR3 amino acid sequences shows come overlapping, but also distinct Kmer usages, amongst CDR3 sequences from vaccine-primed and infection-primed individuals for both the CDR3α (Fig. 3g) and CDR3β (Fig. 3h). VVN and CVV Kmers are more commonly used in CDR3α sequences from vaccine-primed individuals, but are also found in cells from infection-primed individuals at a lower frequency. However, CDR3α sequences from infection-primed individuals show usage of 9 Kmers that are not found in CDR3 sequence from vaccine-primed individuals. Similarly, for CDR3β sequences, cells from vaccine-primed individuals show increased frequency of QPQ, PQH and QHF compared to cells from infection-primed individuals. However, cells from infection-primed individuals show usage of 8 Kmers not found in cells from vaccine-primed individuals.

These differences are reflected in the overall CDR3 motif patterns (Supplemental figure 2h), showing more diverse amino acid usage for CDR3 sequences from infection-primed individuals across CDR3 lengths of 12-15 amino acids, compared CDR3 sequences from vaccine-primed individuals. Comparing the CDR3 lengths, cells from infection-primed individuals show a broader CDR3α length distribution, while cells from vaccine-primed individuals show a more restricted length distribution (Fig. 3i), although both have a peak CDR3α length of 14 amino acids. Comparing the CDR3β length distributions (Fig. 3j), overall the CDR3β lengths from infection-primed individuals appear shorter (14 amino acid length peak), compared to cells from vaccine-primed individuals which appear longer (16 amino acid length peak).

These results suggest that TCR repertoire from infection-primed individuals is more diverse than the vaccine-primed individuals, which appear more restricted. Vaccination may induce a more constrained CDR3α repertoire compared with the more heterogeneous infection-primed repertoires, while CDR3β repertoires may retain greater inter-individual variability and less convergence than the α-chain repertoires, as reflected in their Kmer usages and length distributions.

### 4. Spike-specific CD4^+^ T cells from infection-primed individuals exhibit distinct transcriptomic profiles

To investigate further differences in these spike-specific T cells between the two timepoints, we carried out a transcriptional analysis of the scRNAseq data. Our transcriptomic dataset comprised 2596 total cells collected from the two groups of individuals (infection-primed = 1354 cells, vaccine-primed = 1242 cells) across three epitope specificities (1422 S_166-180_-specific, 607 S_751-765_-specific and 567 S_866-880_-specific cells) from 14 total individuals (9 infection-primed and 5 vaccine-primed), with an average 185±90 cells per individual. Of these cells, a TRA or TRB clonotype was annotated to 67.4% of the cells. All three spike-specific T cells from the different individuals were analysed together.

Unsupervised clustering identified 10 distinct clusters based solely on their gene expression profiles (Fig. 4a-b). A unique cluster of cytotoxic T cells (CD4+_CTL) was identified, clustering independently from the rest of the cell clusters. In addition to cytotoxicity-related genes (e.g. *GNLY, GZMH, GZMA, NKG7* and *GZMB*), these cells also expressed *FGFBP2*, *ZEB2*, *ID2* and *S1PR5*. Other clusters that were identified included a stem-like T memory cluster (CCR7+TCF7+_Stem-like, marker genes include *CCR7*, *NOG*, *LEF1* and *IL6ST*), a SELL+_TCM cluster, a GPR183+_memory cell cluster and a resting memory cluster (Memory_CD4). Furthermore, we identified 5 clusters with evidence of differing levels of activation, including an inflammatory activated cluster (expressing *LCK*), an early activates cluster (expressing *FOS*, *RGCC*, *ANXA1* and *DUSP1*), an acute activated cluster (expressing *DUSP1* and *FOS*), an NKFBIA+ cluster and a SAT1+ cluster. Clusters were generally comprised of cells from all individuals and sequencing runs, indicating the clustering analysis does not represent individual-specific subpopulations or batch effects.

**Fig. 4.**
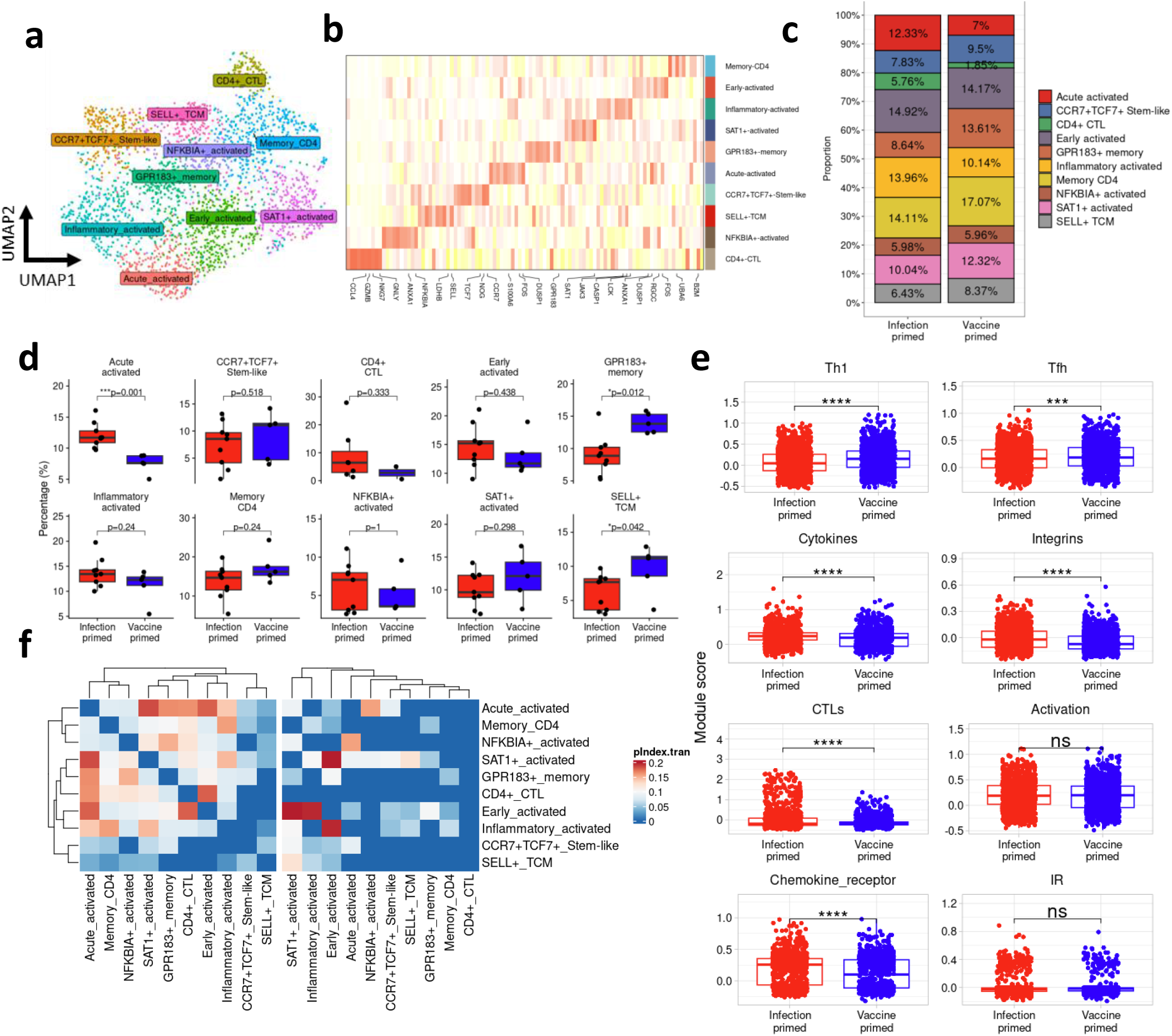
Transcriptomic comparison of cell phenotypes between infection-primed and vaccine-primed individuals. (a) UMAP plot of unsupervised clustering of all spike-specific cells (b) Heatmap of the top 10 marker genes for each cluster, showing scaled expression of the genes across all cells (red=high, yellow=low) (c) Proportion of cells in each cluster, split by whether cells are from infection-primed or vaccine-primed individuals. (d) Boxplots of per individual proportions of cells within each cluster, split by mode of priming, p values are from a linear model corrected for months since last antigen encounter. (e) Per cell module score comparison between two groups, with p-values from a linear model adjusted for months since last antigen-exposure and epitope (f) Paired TCR clonotype sharing between clusters, split by priming (infection primed = left heatmap, vaccine-primed = right heatmap).

We next investigated whether there were any proportional differences in the number of cells in the different clusters originating from individuals primed by infection or vaccination. Acute activated, inflammatory activated and CD4^+^ CTL clusters appeared to contain more cells from individuals who were primed by infection compared to individuals primed by vaccination (Fig. 4c), although on the acute activated cluster was statistically significant (Fig. 4d, p=0.001). On the other hand, GPR183+ memory, SELL+ TCM and SAT1+ clusters appeared to contain more cells from vaccine-primed individuals compared to infection-primed individuals (Fig. 4c), although only the differences within the GPR183+ memory and SELL+ TCM clusters were statistically significant (Fig. 4d, p=0.012 and 0.042 respectively). However, these differences we attenuated when accounting for the months since last antigen exposure before sample collection (vaccination or infection), likely due to the low number of individuals.

To further investigate differences between cells from the different modes of T cell priming, we generated module scores using gene lists compiled from the literature (Supplemental table 3) and compared the module score between groups. Spike-specific cells from individuals primed by vaccination showed a greater Th1 and Tfh gene signatures (Fig. 4e, all p<0.05), consistent with previous reports^10,13^; whereas spike-specific cells from individuals primed by infection showed greater cytokine, integrin, CTL and chemokine receptor gene signatures (Fig. 4e, all p<0.05). There was no statistically significant signature in the activation module score or inhibitory receptor module score between the two groups (p>0.05). Overall, these associations were independent of epitope specificity, showing significance across all three epitope-specific cell subsets (Supplemental figure 3a). Furthermore, sensitivity analyses adjusting for severity and age reveals consistent results independent of disease severity and age of participants (Supplemental figure 4a+b). Investigating the clonal dynamics between clusters, there is more inter-cluster clonotype sharing between cells from infection-primed individuals (Fig. 4f, left panel) compared to cells from vaccine-primed individuals (Fig. 4f, right panel).

Taken together, these results suggest that cells from infection-primed individuals exhibit a more cytotoxic gene profile compared to vaccination-priming, in which cells appear to exhibit more of a Th1 and Tfh gene signature. Cluster clonal dynamics are consistent with the earlier findings of a more constrained TCR repertoire in vaccine-primed individuals, suggesting more phenotype plasticity amongst spike-specific CD4^+^ T cells from infection-primed individuals.

### 5. Increased integrin α4β1-expressing spike-specific CD4^+^ T cells in infection-primed individuals

To investigate the gene expression differences between cells from vaccine-primed individuals compared to cells from infection-primed individuals, to overcome dropout issues associated with scRNAseq data, we pseudo-bulked the gene expression on an individual, epitope and mode of priming level (see Methods for more details). We carried out differential gene expression using limma^23^, while adjusting for epitope and months since last antigen exposure (Fig. 5a). 113 genes showed increased expression in vaccine-primed individuals (p<0.05), including *ITGA4*, *GZMA* and *SELPLG* (Fig. 5b), while 163 genes showed increased expression in infection-primed individuals, including *IFIT2*, *BCL10* and *ADNP*. However, none of these remained significant after multiple testing correction.

**Fig. 5.**
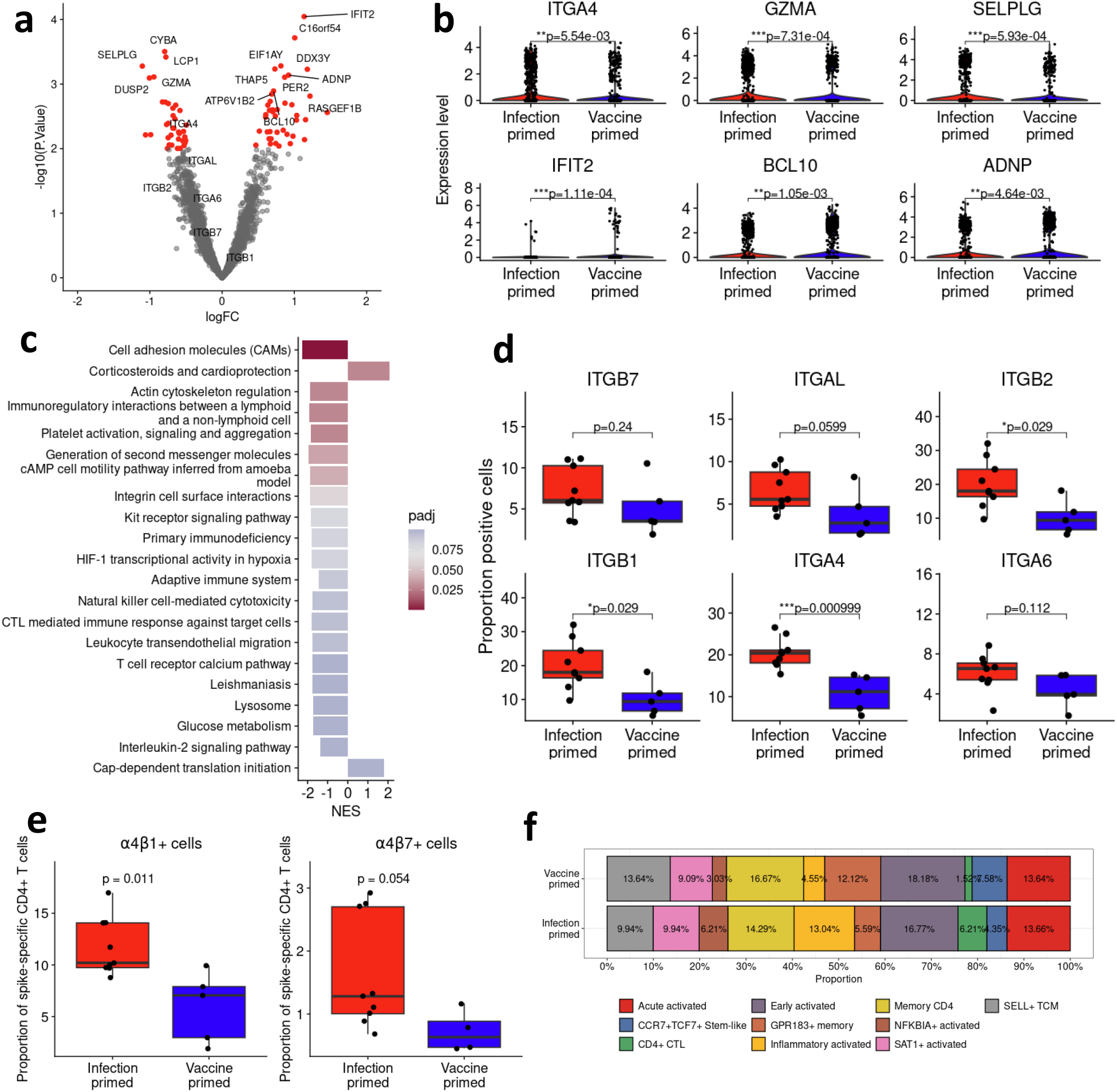
Increased integrin expression and α4β1+ in infection-primed individuals. (a) Pseudo-bulk DEG analysis of vaccine-primed vs infection-primed (positive logFC = higher in vaccine-primed individuals, negative logFC = higher in infection-primed individuals) – red=pvalue <0.01. (b) Violin plot of single-cell expression of select genes, top 3 showing higher expression in infection-primed individuals, lower 3 lower higher expression in vaccine-primed individuals. P-values from analysis in a. (c) GSEA of gene expression analysis between infection-primed and vaccine-primed individuals. Negative NES score (x-axis) = higher in infection-primed. All have adjusted p < 0.1 (d) Proportion of cells per individual with >0 expression of individual integrin subunits at the single-cell level, split by infection-primed (red) and vaccine-primed (blue). (e) Proportion of cells per individual expressing both *ITGA4* and *ITGB1* (α4β1+, left panel) and cells expression both *ITGA4* and *ITGB7* (α4β7+, right panel) cells between the two groups. (f) Phenotype of α4β1+ cells from infection-primed and vaccine-primed individuals. Group wise comparisons in d+e carried out using Wilcoxon signed rank test. NES=normalized enrichment score, logFC=log2 fold change

Given none of the differentially expressed genes remained significant after multiple testing, we carried out gene set enrichment analysis to determine whether there were smaller coordinated changes in gene expression compared to larger changes in individual genes. 21 pathways showed significant enrichment after multiple testing correction (Fig. 5c, adjusted p-value < 0.1), including 19 pathways significantly enriched in infection-primed individuals, and 2 pathways significantly enriched in vaccine-primed individuals. The top significantly enriched pathway in infection-primed individuals is Cell adhesion molecules (CAMs) (NES=-2.25, adjusted p=2.01×10^−4^), with other enriched pathways including Immunoregulatory interactions between a lymphoid and a non-lymphoid cell (NES=-1.90, adjusted p=0.028), Integrin cell surface interactions (NES=-1.86, adjusted p=0.060) and CTL mediated immune response against target cells (NES=-1.78, adjusted p=0.094). Enriched pathways amongst cells from vaccine-primed individuals were Corticosteroids and cardioprotection (NES=2.07, adjusted p=0.028) and Cap-dependent translation initiation (NES=1.79, adjusted p=0.099). The enrichment of integrin-related and cell adhesion-related pathways remain significantly enriched in infection-primed individuals after adjustment for age of participants and disease severity (Supplemental figure 4c+d).

As integrins play a key role in the migration and tissue localisation of T cells^24^, and cell adhesion molecules and integrin-related pathways were amongst the top enriched pathways amongst infection-primed individuals, we investigated the expression of individual integrin molecules between vaccine-primed and infection-primed individuals (Fig. 5d). Infection-primed individuals showed increased expression of *ITGB2* (p=0.029), *ITGB1* (p=0.029) and *ITGA4* (p=0.001), with trends for increased expression of *ITGAL* (p=0.0599), *ITGB7* (p=0.24) and *ITGA6* (p=0.112). ITGA4 forms a heterodimeric complex with ITGB1, forming VLA4 and ITGB7, forming LPAM1. Therefore, we investigated whether there were differences in the proportion of α4β1+ and α4β7+ cells between infection-primed and vaccine-primed individuals (Fig. 5e). Infection-primed individuals showed greater proportion of α4β1+ spike-specific CD4^+^ T cells (p=0.011) and α4β7+ spike-specific CD4^+^ T cells (p=0.054), compared to vaccine-primed individuals, with similar increases in α4β1+ CD4^+^ T cells in infection-primed individuals across all three specificities (Supplemental figure 3b). Moreover, a greater proportion of cells from infection-primed individuals also express αLβ2, although this does not reach statistical significance (p=0.17, Supplemental figure 3c). Furthermore, adjustment for disease severity does not alter the significant increase in α4β1+ spike-specific CD4^+^ T cells in infection-primed individuals, while age-adjustment attenuates the significance slightly (Supplemental figure 4e). In infection-primed individuals, α4β1+ spike-specific cells show a more NFKBIA+ activated, inflammatory activated and CTL phenotype compared to vaccine-primed individuals (Fig. 5f), where α4β1+ cells exhibit a more SELL+ TCM and GPR183+ memory phenotype. Similar phenotypes are seen for α4β7+ spike-specific CD4^+^ T cells between infection-primed and vaccine-primed individuals (Supplemental figure 3d), although only 30 cells were α4β7+.

### 6. Spike-specific CD4+ T cells from infection-primed individuals show greater cytotoxic potential with distinct clonotype dynamics

Given the increased cytotoxicity module score in cells from infection-primed individuals compared to cells from vaccine-primed individuals (Fig. 4e), we next sought to investigate the cytotoxicity-related differences further.

The cytotoxicity module score was comprised of 8 individual genes. The expression of the individual genes shows the score is primarily driven by the expression of *GZMA* and *GNLY*, with low expression of other Granzyme molecules. *GZMA* and *GNLY* were expressed in more cells from infection-primed individuals compared to vaccine-primed individuals (24.59% vs 11.11% respectively for *GZMA* and 10.56% vs 4.27% respectively for *GNLY*), as well as at a higher level in these cells (Fig. 6a). Of the cells that do express the other cytotoxicity-associated molecules, these are expressed in more and at higher levels in infection-primed individuals compared to vaccine-primed individuals (Supplemental figure 5a), including *GZMB*, *NKG7* and *PRF1*, although only <7% of cells in our dataset express cytotoxic molecules other than *GZMA* and *GNLY*. Cells were classed as having a high cytotoxic potential if their cytotoxicity module score was greater than 75^th^ percentile, with 649 cells showing a high cytotoxic potential (Supplemental figure 5b). Of these, 66.26% originate from individuals who were infection-primed compared to only 33.74% from vaccine-primed individuals (Fig. 6b, χ^2^ p=1.49×10^−16^). Cells with no cytotoxic potential showed similar proportions between infection-primed and vaccine-primed individuals (Fig. 6b).

**Fig. 6.**
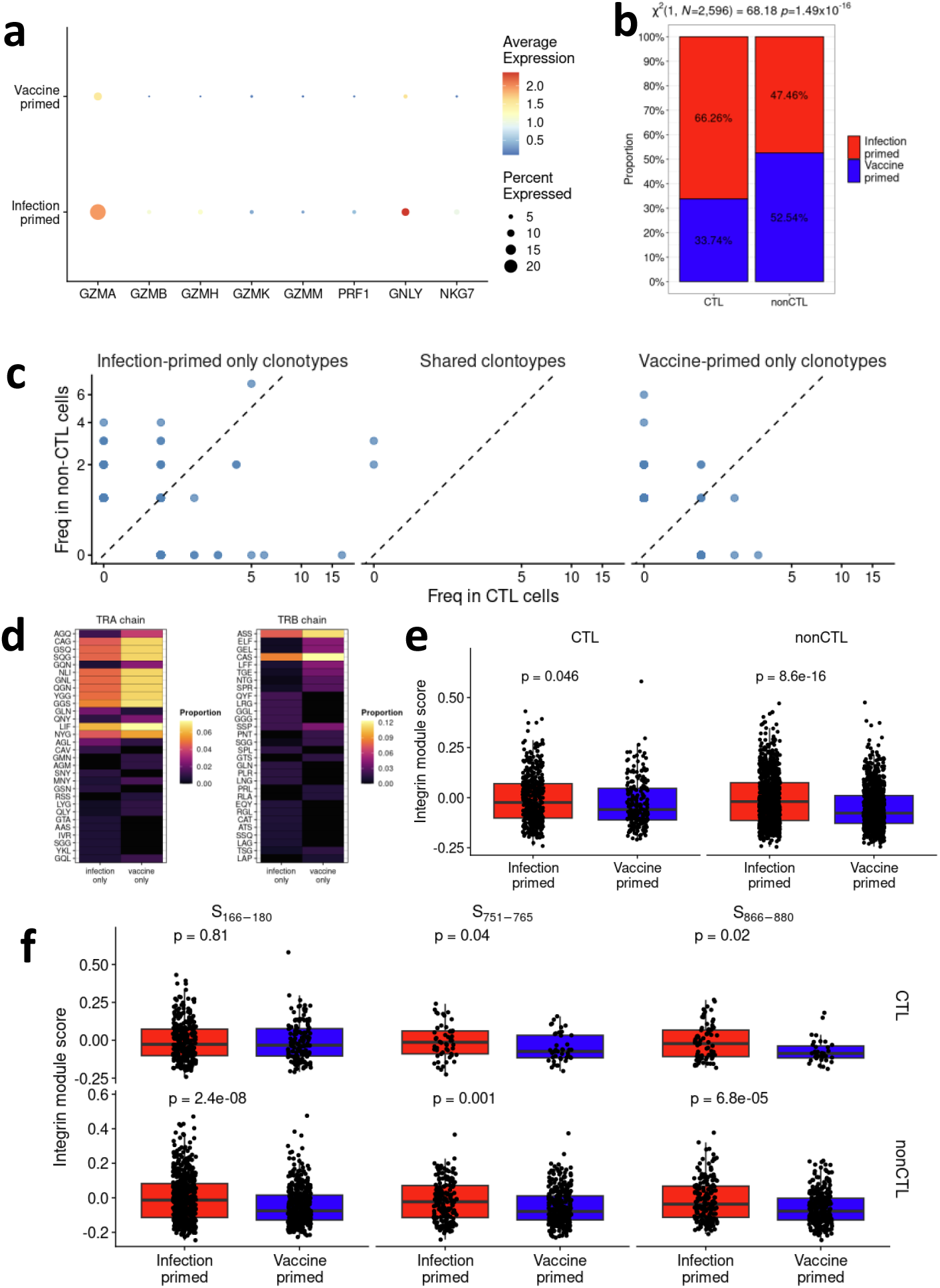
Increased cytotoxic signatures in infection-primed individuals associated with integrin gene signatures. (a) Dotplot of expression of cytotoxicity-related genes between infection-primed and vaccine-primed individuals. (b) Proportion of cells with a CTL module score > 75th percentile between priming. More infection-primed have a higher CTL module score (c) Comparison of the frequencies of the clonotypes with high CTL module scores that found only in infection-primed cells versus in vaccine-primed cells. (d) Differences in CTL clonotypes are reflected by different Kmer usages in CTL cells between infection-primed and vaccine-primed individuals. (e) Comparison of integrin module scores in CTL and non-CTL subset between infection-primed and vaccine-primed individuals. (f) Comparison of integrin module scores in CTL and non-CTL subset within each spike-specific T cell populations between infection-primed and vaccine-primed individuals. χ^2^ test of independence used in b. Wilcoxon signed-rank test used to compare between groups in e and f.

We next investigated the TCR repertoire of the cells with high cytotoxic potential. Cells were classed as having a TCRαβ clonotype found only in infection-primed individuals, found only in vaccine-primed individuals or found in both, and the presence of these clonotypes were compared between cells with high and low cytotoxic potential (Fig. 6c). Only two paired TCRαβ clonotypes were found in both infection-primed and vaccine-primed individuals, and these were found in cells with low cytotoxic potential (Fig. 6c, middle panel). TCRαβ clonotypes found only in infection-primed individuals were found in both cells with high and low cytotoxic potential, although they were found in a greater number of cells with a high cytotoxic potential (Fig. 6c, left panel) compared to TCRαβ clonotypes found only in vaccine-primed individuals (Fig. 6c, right panel).

Moreover, clonotype differences in the cells with high cytotoxic potential are evident from the Kmer usages of their CDR3α and CDR3β sequences (Fig. 6d). Although several Kmers for both the alpha and beta CDR3 sequences exhibit increased proportions in clonotypes found only in vaccine-primed individuals (e.g. LIF and GGS for CDR3α; ASS and SSP for CDR3β), clonotypes found only in infection-primed individuals show increased usage of unique Kmers not found in clonotypes from vaccine-primed individuals, highlighting global repertoire differences in the alpha and beta chains for different modes of priming for spike-specific CD4^+^ T cells.

Consistent with earlier findings, cells with both high and low cytotoxic potential from infection-primed individuals exhibit greater expression of the integrin module score, compared to the cells from vaccine-primed individuals (Fig. 6e, p=0.046 and 8.6×10^−16^ respectively). However, there were differences when comparing different epitope specificities (Fig. 6f). S_751-765_- and S_866-880_-specific cells from infection-primed individuals with both low and high cytotoxic potential exhibited increased integrin module score. However, for S_166-180_-specific CD4^+^ T cells, only cells which low cytotoxic potential from infection-primed individuals exhibited increased integrin module score (p=2.4×10^−8^), whereas cells with a high cytotoxic potential showed no difference in their integrin module score between infection-primed and vaccine-primed individuals (p=0.81).

### 7. *In vitro* cytokine secretion of antigen-specific T cell lines from infection-primed and vaccine-primed individuals

To confirm the findings from the single-cell analysis, we examined the cytokine secretion of S_166-180_- and S_866-880_-specific bulk T cell lines (containing polyclonal T cells) in response to peptide stimulation using Luminex assay from individuals primed by either infection (n=7 individuals) or vaccination (n=6 individuals).

Consistent with increased cytotoxic potential and migratory signals in the spike-specific CD4^+^ T cells from infection-primed individuals, T cell lines stimulated with the S_166-180_ peptide show an increased MCP1 secretion in infection-primed individuals than vaccine-primed individuals overall (Fig. 7a, Type III Wald χ^2^=9.33, p=2.25×10^−3^), and the concentration-response relationship differed between groups (interaction χ^2^=12.86, p=3.35×10⁻⁴). Similarly, we observed an increased IFNγ secretion in infection-primed individuals than vaccine-primed individuals overall (Fig. 7b, Type III Wald χ^2^=12.56, p=3.94×10^−4^), and the concentration-response relationship differed between groups (interaction χ^2^=20.51, p=5.94×10^−6^). These results indicate a steeper increase in both MCP1 and IFNγ production in infection-primed individuals.

**Fig. 7.**
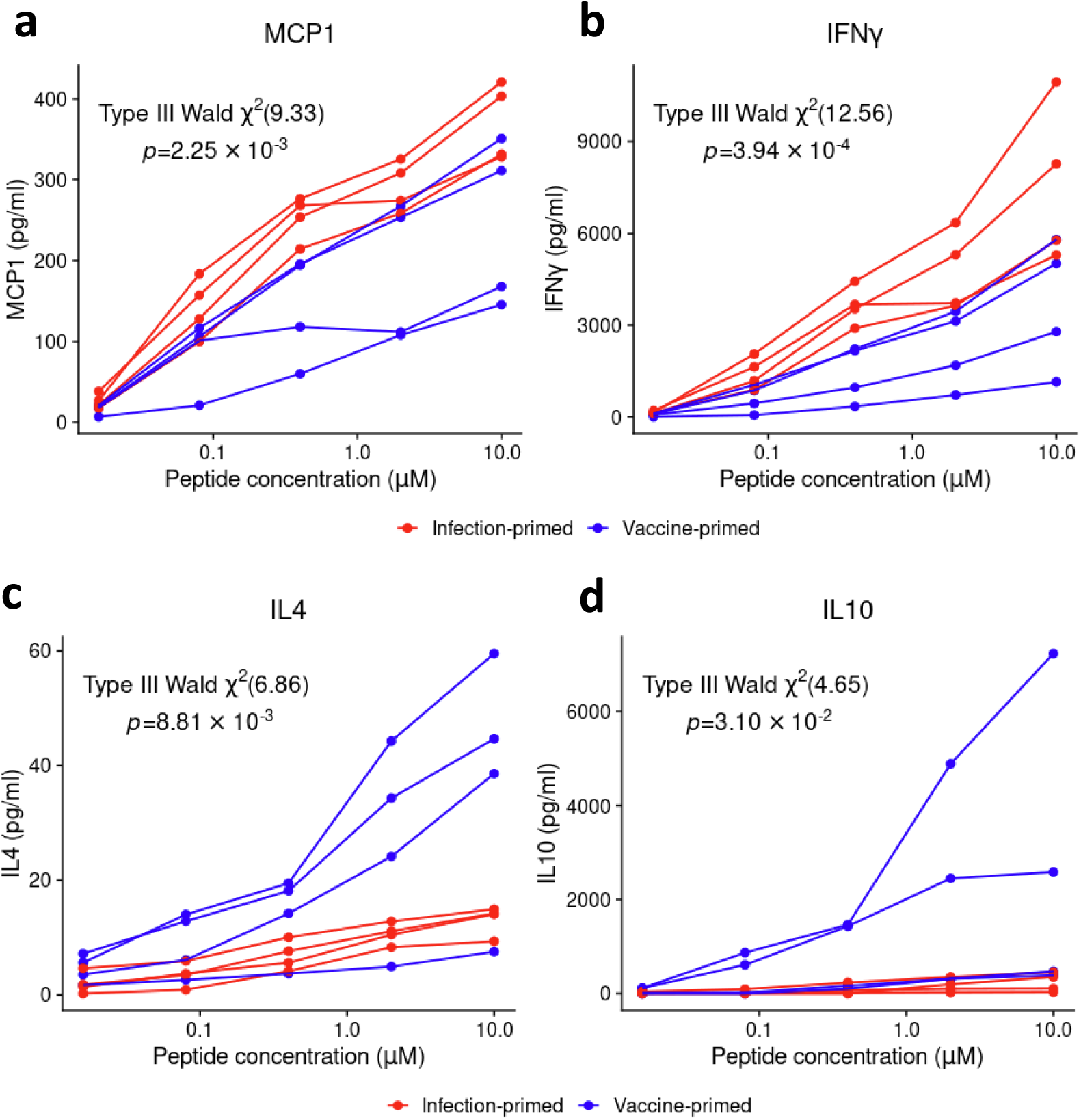
Cytokine secretion by antigen-specific polyclonal T cell lines from infection-primed and vaccine-primed individuals. (a) MCP1, (b) IFNγ, (c) IL4 and (d) IL10 secretion by S_166-180_-specific (a and b) and S_866-880_-specific (c and d) T cell lines were measured by Luminex assay following stimulation with the corresponding peptide. Data from infection-primed individuals are shown in red, and data from vaccine-primed individuals are shown in blue. Group comparisons were carried out using a Type III Wald ANOVA using the following model: priming_mode*log_conc + (1| individual) with priming group p-values reported.

Moreover, consistent with the increased Tfh module score in cells from vaccine-primed individuals and previous reports^13^, S_866-880_-specific T cells stimulated with peptide show an increased secretion of IL4 in vaccine-primed individuals than infection-primed individuals overall (Fig. 7c, Type III Wald χ^2^=6.86, p=8.81×10^−3^), and the concentration-response relationship differed between groups (interaction χ^2^=17.51, p=2.86×10⁻^5^). Surprisingly, T cells from vaccine-primed individuals also show increased secretion of IL10 in response to peptide stimulation (Fig. 7d, Type III Wald χ^2^=4.65, p=3.1×10^−2^), and the concentration-response relationship differed between groups (interaction χ^2^=10.17, p=1.43×10⁻^3^), suggesting a regulatory phenotype of the cells from vaccine-primed individuals.

## Discussion

In this study, we used SARS-CoV-2-specific CD4^+^ T cells as a model to investigate whether the mode of priming influence antigen-specific T cell phenotypes 3-4 years after initial priming. Although HLA-DP typing was not carried out in the vaccination cohort, HLA-DPB1*04:01 has an estimated population frequency of approximately 50%^25^ and we observed an immunodominance of S_166-180_, S_751-765_ and S_866-880_ epitopes in eliciting a CD4^+^ T cell response in vaccine-primed individuals, as has been reported previously in SARS-CoV-2 convalescent individuals^17^. Moreover, the frequency of tetramer+ antigen-specific CD4^+^ T cells recognising these epitopes were comparable between these two group of individuals 3-4 years after the initial priming event. Importantly, these immunodominant spike-specific CD4^+^ T cells analysed in both cohorts were initially primed either by natural SARS-CoV-2 infection or vaccination and subsequently experienced multiple rounds of antigen-exposure through booster vaccinations, reinfections, or both. This unique immunological history provided a valuable opportunely to investigate whether the nature of the initial priming event has long-lasting molecular and functional influences on human memory CD4^+^ T cells despite repeated antigen encounters over several years. By integrating ex vivo single-cell RNA sequencing, paired TCR sequencing and *in vitro* functional analyses, we characterise the transcriptional, clonal and functional profile of Spike-specific CD4^+^ T cells across both these two cohorts, revealing a lasting influence of the priming environment on human CD4^+^ T cell memory with differences in TCR repertoire organisation, cytotoxic potential, and integrin-associated migratory programmes.

Our single-cell RNA-sequencing analysis of tetramer-sorted S_166-180_, S_751-765_ and S_866-880-_specicfic CD4^+^ T cells revealed that these immunodominant spike-specific CD4^+^ T cells exhibited distinct transcriptional signatures and functional properties according to the mode of priming. Infection-primed cells exhibited a more effector-like transcriptional profile characterised by increased cytotoxicity-, cytokine- and chemokine-associated signatures, whereas vaccine-primed cells showed enrichment of Tfh- and central memory-associated programmes. These findings are broadly consistent with previous studies^13,17^. Germinal centre (GC) responses occur in the draining lymph nodes, and play a critical role in developing long-lasting antibody responses^26^. Tfh cells in the lymph node are necessary for both forming and sustaining the GC and formation of long-lived plasma cells and memory B cells. Mudd et al.^13^ show that Tfh cell responses are in key establishing long-term immunity after vaccination. Our results suggest that vaccine-priming in the lymph nodes may result in a long term Tfh-like phenotype in antigen-specific CD4^+^ T cells that may play a role in the long-term T cell memory and B cell help in the GCs, compared to infection-priming.

Furthermore, in antigen-specific CD4^+^ T cells from infection-primed individuals, we identified an enrichment of cell adhesion molecule and integrin-associated pathways. We identified increased expression of multiple integrin subunits and a greater frequency of α4β1-expressing and αLβ2-expressing spike-specific CD4^+^ T cells in infection-primed individuals. Integrins are heterodimeric cell surface molecules that play fundamental roles in cell interactions. In T cells, integrins play key roles in lymphocyte trafficking and tissue retention, T cell activation, proliferation and differentiation. Although α4β1 and αLβ2 are involved in T cell migration^27–29^, they also play key roles in the immune synapse^30,31^, acting as costimulatory molecules and enhancing T cell activation and differentiation, and reorganisation at the immune synapse in an antigen-dependent manner. Mittelbrunn et al.^30^ show that the engagement of α4 integrins promotes CD4^+^ T cells towards a Th1 polarization, resulting in a secretion of IFNγ and inhibiting IL4 secretion, consistent with our cytokine secretion results from infection-primed and vaccine-primed individuals after peptide stimulation. Furthermore, α4β1-expressing cells from infection-primed individuals were associated with inflammatory and cytotoxic transcriptional profiles. These findings also raise the possibility that distinct priming environments may establish long-lasting migratory and effector functions that persist within circulating memory T cell populations.

Using an AIMs assay to identify spike-specific CD4^+^ T cells, Gray-Gaillard et al.^32^ identify a robust enrichment for interferon signalling and inflammatory responses in spike-specific CD4^+^ T cells from infection-primed individuals both pre- and post-vaccine boosters, compared to vaccine-primed individuals. Given the lack of change resulting from booster immunization, this suggests that the early antigen exposure results in a phenotype in the cells that is maintained regardless of subsequent antigen exposure. The authors extend this to show that epigenetic changes as a result of inflammation was associated with infection-priming leading to cytotoxic profile of spike-specific cells. However, these epigenetic changes are occurring after AIMs activation, and T cell activation has been shown to elicit widespread epigenetic changes from their basal state^33,34^. In this study, we show that this inflammatory-enriched phenotype and cytotoxic potential is present in spike-specific CD4^+^ T cells ex vivo, 3-4 years after initial antigen exposure in infection-primed individuals, accompanied by an increased migratory potential mediated by α4β1 integrins. Although not possible in our dataset, epigenetic changes, which are known to persist long-term, may contribute to the integrin and cytotoxic signatures seen in infection-primed individuals, leading to the increased expression of α4β1 and αLβ2 integrins. Furthermore, while the increased frequencies of αLβ2+ cells in infection-primed individuals did not reach significance, likely due to low numbers of individuals, the differences observed suggests that infection priming may result in a selective reprogramming of specific trafficking and differentiation pathways within antigen-specific CD4^+^ T cells that is not fully recapitulated by vaccination. A larger pool of α4β1+ memory CD4^+^ T cells may therefore lead to a more rapid recruitment to infected respiratory tissue following re-exposure. These findings warrant further investigation into the functional validation of α4β1 and αLβ2 integrins in infection-primed and vaccine-primed individuals and their role in the function of the long-term memory T cell population. Although the role of these integrins in T cell migration is well-established, it remains important to determine the mechanisms contributing to the upregulation of these integrins in the infection-primed memory CD4^+^ T cell population and any further roles they may play in T cell migration and signalling, allowing for better vaccine development that may promote the formation of a memory pool with increased migration potential.

Consistent with this, functional validation experiments demonstrated enhanced IFNγ and MCP-1 production by infection-primed CD4^+^ T cells following peptide stimulation, supporting the enhanced effector and cytotoxic transcriptional signatures observed at the single-cell level. Monocyte chemoattractant protein-1 (MCP-1/CCL2) is an inflammatory cytokine that recruits monocytes and macrophages to sites of inflammation^35^. Increased MCP-1 secretion by antigen-specific CD4^+^ T cells from infection-primed individuals suggests an increased capacity to recruit monocytes and other innate immune cells to sites of inflammation. Taken together, these findings suggest that infection-priming may generate long-term memory CD4^+^ T cells that are both functionally poised and equipped to coordinate local immune responses within peripheral tissues.

Although we cannot be certain, one contributing factor to these long-term differences may be the site of priming. Previous literature suggests that the microenvironment surrounding T cells during initial antigen exposure can influence T cell activation and differentiation trajectories^1,2,6,8^. SARS-CoV-2 viral infection initiates immune responses within the respiratory tract, where antigen is encountered in the context of local inflammation and tissue-resident antigen-presenting cells. In contrast, vaccination primarily induces immune responses within draining lymph nodes following antigen transport from the site of injection^36^. Distinct antigen-presenting cell populations, cytokine environments and tissue-derived signals encountered during priming are known to influence both T cell differentiation and homing receptor expression, resulting in differential imprinting by tissue-specific signals present during respiratory infection but absent during vaccine-mediated priming. However, further functional and mechanistic studies are required to determine whether different APC populations or the different environmental milieu contribute to the phenotype of long-term antigen-specific CD4^+^ memory cell populations.

Beyond the phenotypic and functional differences, our single-cell paired TCR repertories analysis revealed subtle but consistent differences in these immunodominant spike-specific CD4^+^ T-cell repertoire associated with the mode of priming. Consistent with our previous report^17^, we identified the same public TCR clonotypes in vaccine-primed individuals as those previously observed in infection-primed individuals, indicating that these public clonotypes can be primed by both natural infection and also vaccination. The presence of these shared public clonotypes suggests the existence of common precursor T-cell populations that are readily recruited into SARS-CoV-2-specific response irrespective of the priming modality; however, this may not necessarily demonstrate mode of priming-independent recognition by T cells, given the relatively high probability of generation of public clonotypes to immunodominant epitopes. Given the reported association of these public T cell clonotype with protective antiviral immunity, the long-term persistence of T cells bearing these receptors may contribute to protection against future infection regardless how they have been primed.

We also identified greater TCR repertoire diversity in infection-primed individuals. Although many clonotypes were shared between the two groups of individuals, infection-primed individuals displayed broader TCR sequence usage and less CDR3α and CDR3β sequence convergence than vaccine-primed individuals, particularly for S_751-765_-specific cells. This was also evident for S_866-880_-specific cells, albeit to a lower extent. Greater sequence convergence is generally thought to reflect preferential recruitment of TCRs with similar structural solutions for peptide-MHC recognition^37^. However, this finding was not evident for S_166-180_-specific TCR repertoires, with a trend for increased diversity in vaccine-primed individuals. This may not be too surprising given the different HLA restriction of the three epitopes, with S_166-180_ restricted by the HLA-DPB1*04:01 allele while the other two epitopes are both restricted by the HLA-DRB1*15:01 allele. Previous studies have highlighted a contribution of HLA-restriction on the TCR repertoire composition^38,39^, which may contribute to this difference and merits further investigation in the context of T cell priming. Despite this, our findings suggest that natural infection may recruit a greater range of TCR clonotypes into the long-term memory pool than vaccination. Several factors could influence this, including prolonged antigen exposure, the heterogenous antigen load associated with natural infection, greater diversity of antigen-presenting cell subsets and distinct inflammatory signals during infection. In contrast, vaccination generally results in a more spatially restricted antigen exposure^9,36^, which may favour the expansion of fewer high-affinity clonotypes. Recent studies have shown several factors that can influence the breadth of the responding TCR repertoire and between clonal expansion and recruitment of rare clonotypes^40–47^. These observations support the concept that the distinct immunological environments generated by infection and vaccination shape not only the magnitude and phenotype of memory CD4^+^ T cell responses, but also the TCR repertoire.

Several limitations should be acknowledged. Our overall study cohort size is relatively small, limiting statistical power for some analyses and preventing assessment of additional clinical variables that may influence memory formation. Furthermore, all vaccine-primed individuals, as well as infection-primed individuals, likely subsequently experienced breakthrough infection and further vaccinations, meaning that the observed differences likely reflect the influence of initial priming rather than exclusive exposure histories. However, we also cannot conclude that the mode of priming is the causal factor in these differences observed 3-4 years after initial antigen-exposure, requiring further mechanistic investigation into the causal links. Despite multiple subsequent antigen encounters in both cohorts, signatures associated with the initial T cell priming remained detectable 3-4 years later. Although we accounted for months since last antigen exposure in most analyses, further replication and validation in larger cohorts remain necessary. Finally, although we report consistent and robust differences in the antigen-specific CD4^+^ T cells from infection-primed and vaccine-primed individuals, future studies using larger cohorts, tissue samples and epigenetic profiling would be required to determine the mechanisms responsible for these persistent transcriptional and repertoire differences.

In conclusion, our findings demonstrate that the mode of priming exerts a durable influence on the composition and phenotype of antigen-specific CD4^+^ T cell memory, 3-4 years after initial priming. Collectively, these results suggest that the first encounter with an antigen, and the subsequent T cell priming event, do not generate T cell responses that only differ in magnitude and breadth, but establishes long-lasting influences on the CD4^+^ T cell memory beyond just the level of response, but also influences the TCR repertoire, migratory potential and effector function years after priming, likely to influence the recall response. Understanding how priming conditions shape these long-lived memory populations may help to inform the design of newer vaccines in the future, against known and novel pathogens, aimed at eliciting specific functional properties of protective T cell immunity.

## Acknowledgments

We are grateful to all the participants for donating their samples and data for these analyses, and the research teams involved in the consenting, recruitment, and sampling of these participants.

This work was supported by the Chinese Academy of Medical Sciences (CAMS) Innovation Fund for Medical Science (CIFMS), China (grant number: 2024-I2M-2-001-1) (T.D., D.J., Y.P., X.Y., G.L., E.A., T.X., J.C.K.). The study is also funded by the UK Medical Research Council (MRC) grant MR/Y015347/1 and MR/R022011/1, IMMPROVE - MR/Y004450/1; China Scholarship Council (G.L); the NIHR Oxford Biomedical Research Centre (J.C.K), Wellcome Trust Investigator Award (204969/Z/16/Z) (J.C.K), Wellcome Trust Grants (090532/Z/09/Z and 203141/Z/16/Z) to core facilities Wellcome Centre for Human Genetics.

This work uses data provided by patients and collected by the NHS as part of their care and support #DataSavesLives.

NIH Tetramer Facility provided S_166-180_(CTFEYVSQPFLMDLE)-DPB1*04:01 monomer.

## Author contributions

T.D. conceptualized the project; T.D. and Y.P. designed and supervised T cell experiments; J.C.K. supervised bioinformatic analysis, A.M. supervised sample collection; G.L., Y.P., X.Y., D.J. performed all T cell experiments and data analysis; E.A. performed single cell data analysis; T.R. performed HLA typing and next generation sequencing; J.W.F. provided MHC Class II Tetramers; D.J. established the heathy donor vaccination cohort and collected blood samples and data. J.C.K, A.J.M, A.F established convalescent cohorts and collected clinical samples and data; C.W., K.C., P.S., and T.X. provided technical assistance and critical reagents; T.D, J.C.K. and Y.P. supervised data analysis, E.A., T.D. and Y.P. wrote the original draft. All authors reviewed and edited the final manuscript and Figures.

## Data and materials availability

All data are available in the main text or the supplementary materials. All other data can be made available by reasonable request to the corresponding author.

**Supplemental Figure 1:**

(a) Flow diagram of initial cohorts, HLA typing of individuals, individuals with positive IFNγ ELISpot results and final selection of individuals for scRNAseq analysis. (b) Representative gating strategy for identification of tetramer+ CD4+ T cells. (c) Gating strategy of index-sorted tetramer+ CD4+ T cells. The specificity and purity of the sorted tetramer+ CD4+ cells used for scRNAseq (red) can be visualized in the overlaid plot relative to the total CD4^+^ population (blue).

**Supplemental Figure 2:**

(a) PCA of TRBV gene usage across S_751-765_-specific cells, coloured by mode of priming (red=infection-primed, blue=vaccine-primed) (b) Box and whisker plot of PC1 values, grouped by mode of priming, showing median±IQR. P-value from a Wilcoxon signed rank test (c) Proportion of TRNV24-1+ cells (blue) or other TRBV usages (red) in cells from infection-primed and vaccine-primed individuals; χ^2^ test of independence to compare between groups. (d) MDS plot of Jensen-Shannon divergence of S_866-880_-specific CDR3α sequences (e) MDS plot of Jensen-Shannon divergence of S_166-180_-specific CDR3α sequences (f) MDS plot of Jensen-Shannon divergence of S_866-880_-specific CDR3β sequences (g) MDS plot of Jensen-Shannon divergence of S_166-180_-specific CDR3β sequences (h) CDR3α sequence alignments for infection-primed (top) and vaccine-primed (bottom) individuals, for sequences 12 amino-acids to 15 amino-acids (left to right).

**Supplemental Figure 3:**

(a) Per cell module score comparison between two groups, split by epitope specificity, with p-values from a linear model adjusted for months since last antigen-exposure and epitope (b) Proportion of cells per individual expressing both *ITGA4* and *ITGB1*, split by epitope specificity. (c) Proportion of cells per individual expressing both *ITGAL* and *ITGB2*. (d) Phenotype of α4β7+ cells from infection-primed and vaccine-primed individuals. P-value from a Wilcoxon signed rank test. P-value from a Wilcoxon signed rank test in b+c.

**Supplemental Figure 4:**

(a) Per cell module score comparison between the two groups, with p-values from a linear model adjusted for months since last antigen-exposure, epitope and age of participants. (b) Per cell module score comparison between the two groups, with p-values from a linear model adjusted for months since last antigen-exposure, epitope and disease severity. (c) GSEA of gene expression analysis between infection-primed and vaccine-primed individuals after additional adjustment for age of participant. (d) GSEA of gene expression analysis between infection-primed and vaccine-primed individuals after additional adjustment for disease severity. For both (c) and (d), negative NES score (x-axis) = higher in infection-primed and adjusted p < 0.1. (e) Proportion of cells per individual expressing both *ITGA4* and *ITGB1* (α4β1+) between the two groups with p-values from a linear model after additional adjustment for age of participants (left) and disease severity (right).

**Supplemental Figure 5:**

(a) Dotplot of expression of cytotoxicity-related genes between infection-primed and vaccine-primed individuals, after removal of *GNLY* and *GZMA*. (b) Per cell CTL module score of all cells, with horizontal line denoting the 75^th^ percentile, used to classify cells as CTL with a module score greater than the horizontal line.

## Methods

### Study participants

#### Convalescent cohort

Participants were recruited from the John Radcliffe Hospital in Oxford, UK, between March 2020 and September 2021 by identification of patients hospitalized during the SARS-CoV-2 pandemic and recruited into the Sepsis Immunomics study. All participants were sampled at least 28 days after symptom onset during the primary infection. A subset of 10 individuals were subsequently sampled around 3-4 years after their initial infection. Written informed consent was obtained from all participants. Ethical approval was given by the South Central-Oxford C Research Ethics Committee in England (ref. 19/SC/0296). Clinical definitions were defined as previously described^48^.

#### Healthy vaccination cohort

68 healthy volunteers were recruited in Oxford, UK, from April 2021. All participants received a primary course of SARS-CoV-2 vaccination with either the ChAdOx1 nCoV-19 (AstraZeneca) vaccine or an mRNA vaccine (BNT162b2 [Pfizer–BioNTech] or mRNA-1273 [Moderna]) before any documented SARS-CoV-2 infection. Peripheral blood samples were collected longitudinally following each vaccine dose and any SARS-CoV-2 infection. SARS-CoV-2 infection was either confirmed by a positive SARS-CoV-2 lateral flow test or, where virological confirmation was unavailable, by the detection of anti-Spike antibodies and/or T cell responses to SARS-CoV-2 membrane (M) and nucleoprotein (N) antigens measured by IFN-γ ELISpot. Five participants were further sampled in 2024, at least 6 months after their most recent SARS-CoV-2 infection or vaccination. Written informed consent was obtained from all participants. Ethical approval was given by London-Bloomsbury Research Ethics Committee in England (ref. 21/PR/1155).

### Enzyme-linked Immunosorbant spot (ELISpot) assay

Ex vivo ELISpot was carried out using either freshly isolated or cryopreserved PBMCs as described previously^11^. Peptides were added to 2 × 10^5^ PBMCs at a final concentration of 2 μM for 16–18 h. To quantify antigen-specific responses, mean spots of the control wells were subtracted from the sample wells, and the results were expressed as spot-forming units (SFU) per 10^6^ PBMCs. Responses were considered positive if results were at least three times the mean of the negative control wells and >25 SFU/10^6^ PBMCs. If negative control wells had >30 SFU/10^6^ PBMCs or positive control wells (Phytohemagglutinin stimulation) were negative, the results were excluded from further analysis.

### Generation of antigen-specific polyclonal T cell lines

Short-term SARS-CoV-2-specific T cell lines were generated as described previously^49^. Briefly, 2 × 10^6^ PBMCs were stimulated with 10 μM peptides at 37 °C for 1 h and cultured in H10 (RPMI 1640 medium with 10% human serum, 2 mM glutamine, 100 units/ml of penicillin and 100 μg/ml of streptomycin) at 2 × 10^6^ cells per well in a 24-well plate (Costar). IL-2 was added to a final concentration of 100 IU/ml on day 3. S_166-180_-, S_751–765_- and S_866–880_-specific bulk polyclonal T cell lines were established by sorting tetramer^+^ CD4^+^ T cells from PBMCs or short-term T cell lines on day 10-14. Bulk T cell lines containing multiple clonotypes were then expanded with irradiated allogeneic PBMCs every 2–3 weeks as described previously^50^

### Cytokine production assessment

To assess cytokine production, *in vitro* expanded antigen-specific bulk T cells were co-cultured with BCLs loaded with or without peptide at an E:T ratio of 2:1. After 48 h, 50 μl of supernatant was collected for cytokine detection. Cytokines, including IFN-γ, TNF-α, IL-2, IL-4, IL-6, IL-10, IL-13, RANTES and GM-CSF were quantified using the Bio-Plex Pro Human Cytokine Assay (Bio-Rad) following the manufacturer’s instructions, then ran on Bio-Plex 200 (Bio-Rad). Concentration was analysed by the Bio-Plex Manager (Bio-Rad).

### Tetramer-associated magnet enrichment and cell sorting of spike epitope-specific CD4^+^ T cells

S_166-180_-, S_751-765_- and S_866-880_-specific CD4^+^ T cells were enriched from 3-4-year samples prior to sorting for scRNA-seq, as previously described^18,19^. In brief, 1.5-3×10^7^ PBMCs were labelled with APC- or PE-conjugated peptide-MHC-class II tetramers (S_166-180_ tetramer from NIH Tetramer Core Facility, S_751-765_- and S_866-880_ tetramers from ProImmune) for 30 minutes. Enrichment was then performed with anti-APC or anti-PE microbeads using magnetic-activated cell sorting technology (Miltenyi Biotec) following the manufacturer’s instructions. Subsequently, enriched S_166-180_-, S_751-765_- and S_866-880_-specific CD4^+^ T cells were stained with CD3-BV786 and CD8-BV510 (BioLegend), CD4-FITC, CD14-PE-CF594, CD19-PE-CF594 and CD16-PE-CF594 (BD Biosciences). Before sorting, cells were stained with Propidium Iodide (PI) (eBioscience) to exclude nonviable cells. CD3^+^CD8^−^CD4^+^tetramer^+^ were sorted for scRNA-seq using a BD FACSAria Fusion sorter or BD FACS Aria III (BD Biosciences).

### SmartSeq2 scRNA-seq

ScRNA-seq of *ex vivo* sorted tetramer^+^ cells was performed using SmartSeq2 analysis as described previously^51^. Reverse-transcription (RT) and PCR amplification were performed with the exception of using ISPCR primer with biotin tagged at the 5′ end and increasing the number of cycles to 25. Sequencing libraries were prepared using the Nextera XT Library Preparation Kit (Illumina) and sequencing was performed on Illumina NextSeq sequencing platform with NextSeq Control Software v.4.

### Single-cell RNA sequencing analysis

BCL files were converted to FASTQ format using bcl2fastq v2.20.0.422 (Illumina). FASTQ files were aligned to human genome hg19 using STAR v2.6.1d. Reads were counted using featureCounts (subread v2.0.0). The resulting counts matrix was analyzed in R v4.0.1 using Seurat v4.0.1. Cells were filtered using the following criteria: minimum number of cells expressing specific gene=3, minimum number of genes expressed by cell=200 and maximum number of genes expressed by cell=4000. Cells were excluded if they expressed more than 10% mitochondrial genes. SCTransform was carried out to normalise and transform the data, regressing out percent of reads aligning to mitochondrial genes, cell cycle stage, epitope specificity and individual, using the ‘vars.to.regress’ parameter. No integration of the data was carried out as no residual batch effect was observed. FindNeighbours and RunUMAP were carried out using 10 PCs and clustering performed using the Leiden algorithm using the igraph method. The FindMarkers function was used to evaluate differentially expressed genes (DEGs) between clusters and cell subsets annotated based on manual curation of the cluster-specific DEGs. The AddModuleScore function was used to calculate a module score for each cell, to investigate the expression of specific gene sets (Supplementary Table 3). Gene sets were manually curated from the literature and filtered to contain only genes present in the single-cell RNAseq dataset. Higher scores indicate that that specific signature is more highly expressed in a particular cell compared with the rest of the population.

### Infection-primed vs vaccine-primed gene expression comparison

For comparison of gene expression between infection-primed and vaccine-primed individuals, the gene expression values were pseudo-bulked using the AggregateExpression function, grouping cells by timepoint, individual and epitope specificity. Gene expression differences between modes of priming was carried out using limma^23^, with the following model design: ~mode + epitope + months_since, adjusting the analysis for epitope specificity and months since last antigen exposure. Gene set enrichment analysis (GSEA) was carried out using the ranked genes from the differential expression analysis, ranking genes based on their log2 fold changes (logFC), with greatest positive logFC at the top of the list (greatest expression in vaccine-primed individuals), and greatest negative logFC at the bottom of the list (greatest expression in infection-primed individuals). GSEA was carried out using gene sets curated from the NCATS BioPlanet 2019 database^52^, and p-values corrected for multiple testing using the Benjamini-Hochberg adjustment.

### TCR sequence reconstruction

TCR sequences were reconstructed from scRNA-seq FASTQ files using MiXCR v.3.0.13^53^ to produce separate TRA and TRB output files for analysis. MiXCR was run using the mixcr analyze shotgun method with the following parameters: -s hsa --starting-material rna --contig-assembly --only-productive --receptor-type tcr. Resulting TCRs were parsed into R and matched to the gene expression data using their cell IDs.

#### TCR repertoire analysis

TCRs were filtered to retain 1/2α or 1β; paired αβ cells consist of 1α1β or 2α1β. Clonotypes were defined as α (CDR3α amino acid + TRAV), β (CDRβ amino acid + TRBV) or paired αβ (CDR3α amino acid + TRAV + CDRβ amino acid + TRBV). TCR repertoire diversity was calculated using the Shannon diversity metric, calculated using the vegan R package. Clonotype sharing plots between individuals were generated using ggalluvial. For epitope-specific analysis, samples were restricted to epitope-specific TCRs with productive CDR3 sequences. For each group CDR3 clonotype frequencies were calculated and normalized to sum to one within each group. Pairwise Jensen–Shannon divergences between these normalized clonotype distributions were calculated and the resulting distance matrix was projected into two dimensions by classical multidimensional scaling (MDS, k = 2). Kmer usage and CDR3 length distributions were analysed using scRepertoire^54^ (v2.6.2). TCR sharing between clusters was calculated using STARTRAC^55^ and transition index visualised as a heatmap for infection-primed and vaccine-primed individuals.

### Statistical analysis

All statistical analyses were carried out in R v4.5.3. Chi-squared test of independence was used to compare ratio difference between two groups. Wilcoxon signed-rank test was employed to compare two groups. Statistical significance was set at p<0.05 and all tests were 2-tailed. A Type III Wald ANOVA was used to compare cytokine secretion between infection-primed and vaccine-primed individuals from the cytokine secretion assay, adjusting the model for repeated measures per individual using the following model design: priming_mode*log(conc) + (1|individual) to investigate both cytokine secretion between groups and whether there were any differences in the slope of the curve between groups (interaction term). MAST test was used to find DEGs between two conditions and represented by volcano plots and violin plots. P values were adjusted for multiple testing using the Benjamini-Hochberg method. (ns, not significant; *p<0.05, **p<0.01, ***p<0.001, ****p<0.0001).

