## Supplementary figures and images for "Mode of T cell priming durably shapes the TCR repertoire, effector function and α4β1 integrin expression of human virus-specific CD4^+^ T cells"

### Supplemental Figure 1

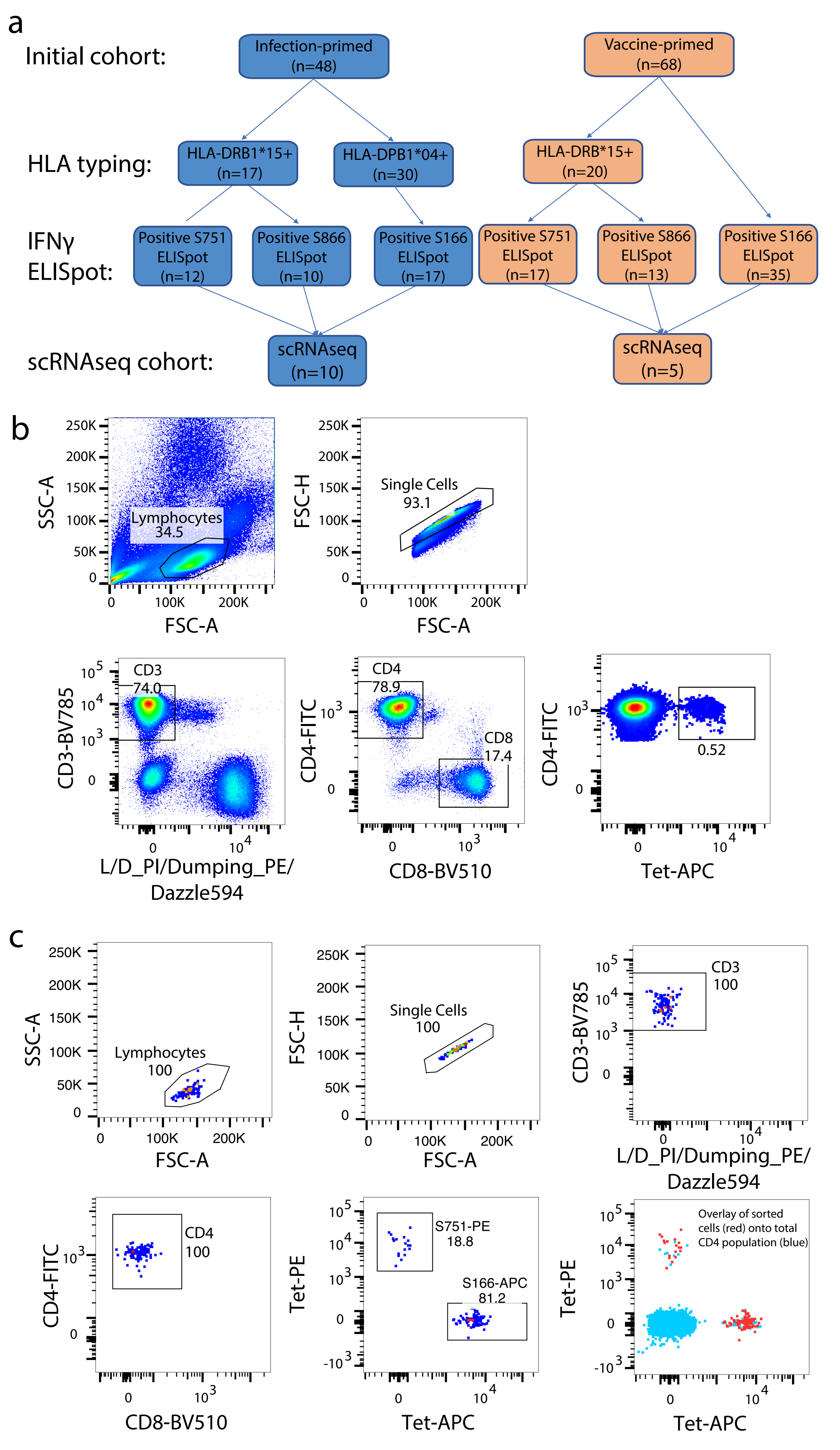

### Supplemental Figure 2

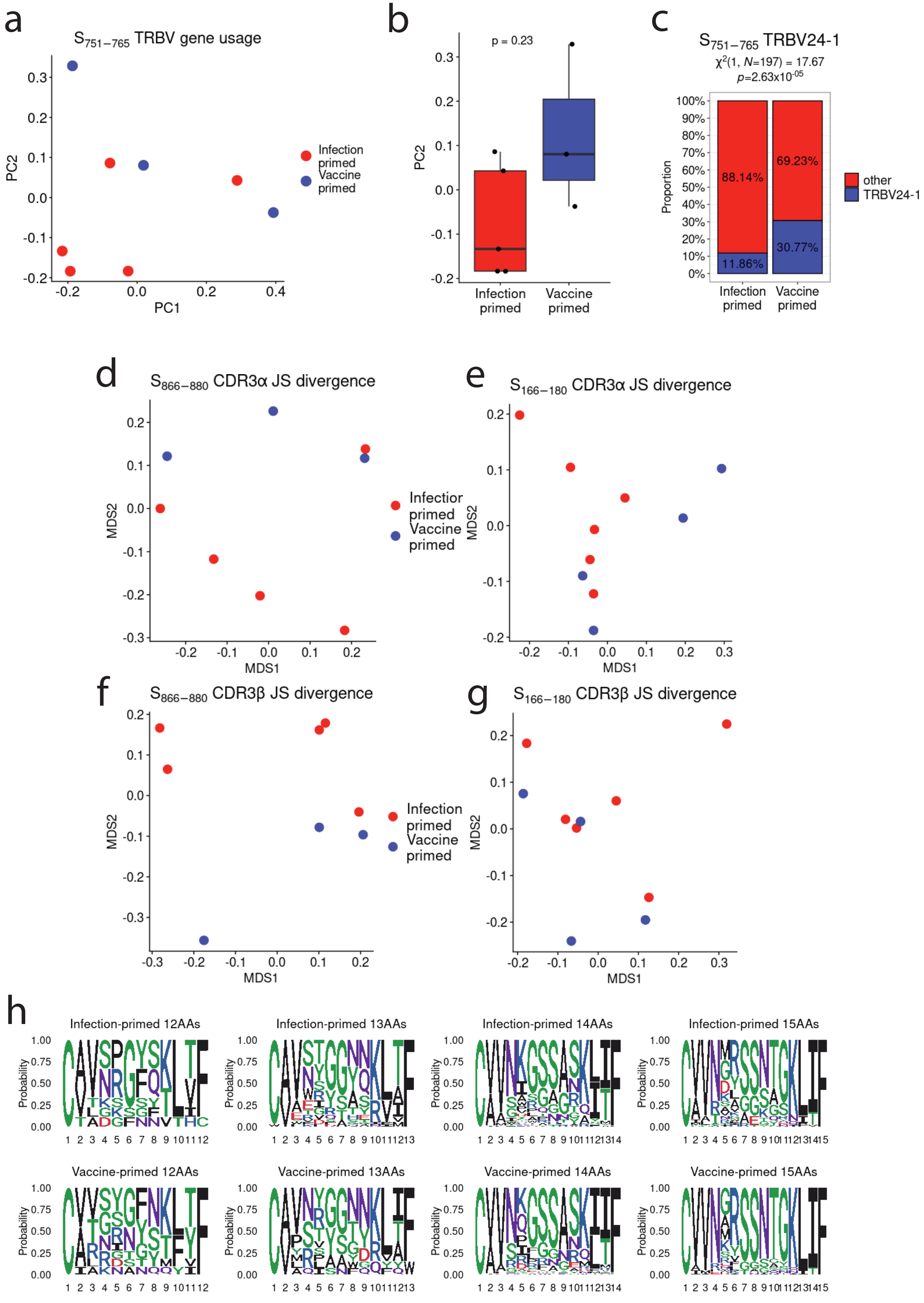

### Supplemental Figure 3

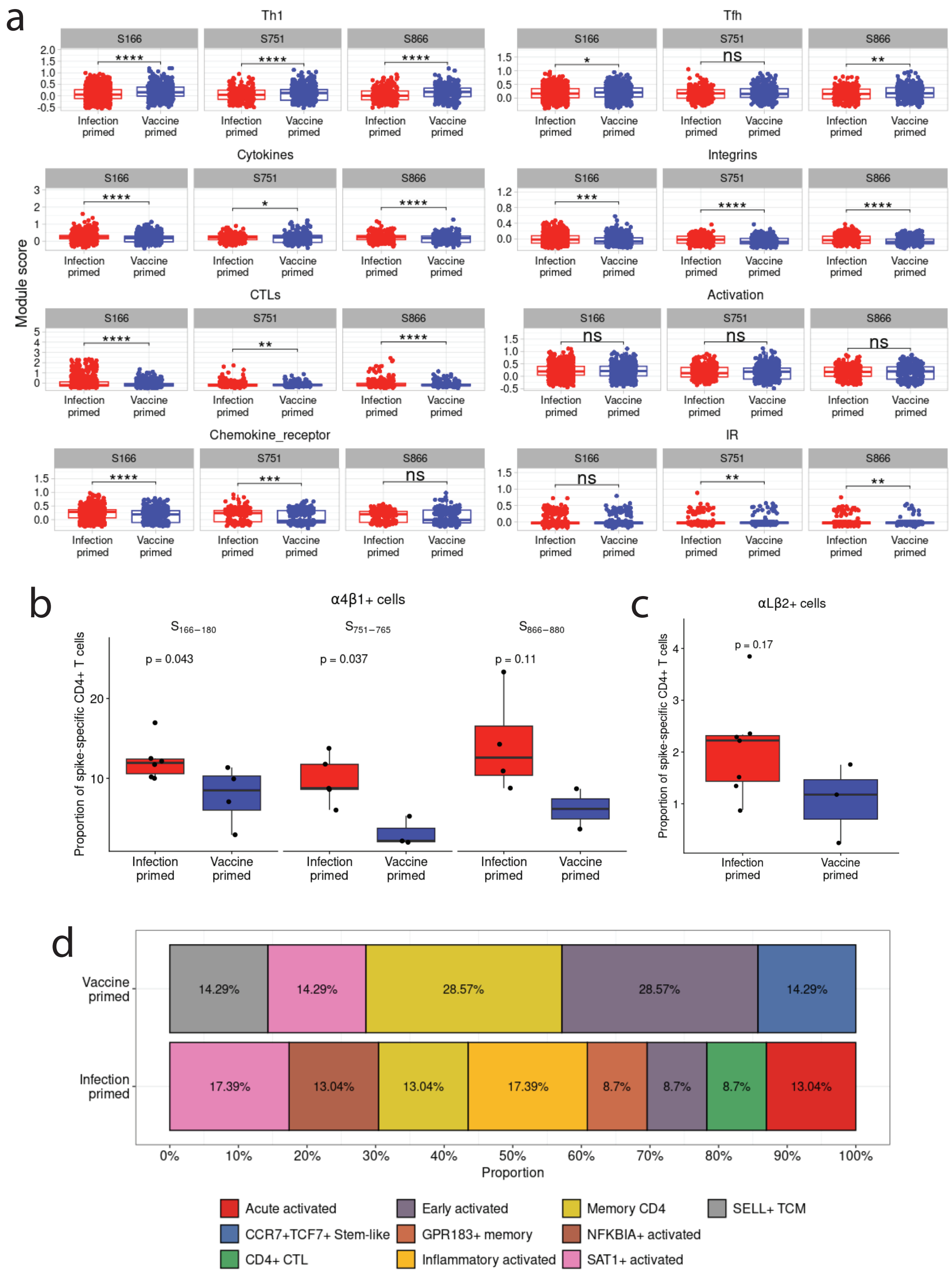

### Supplemental Figure 4

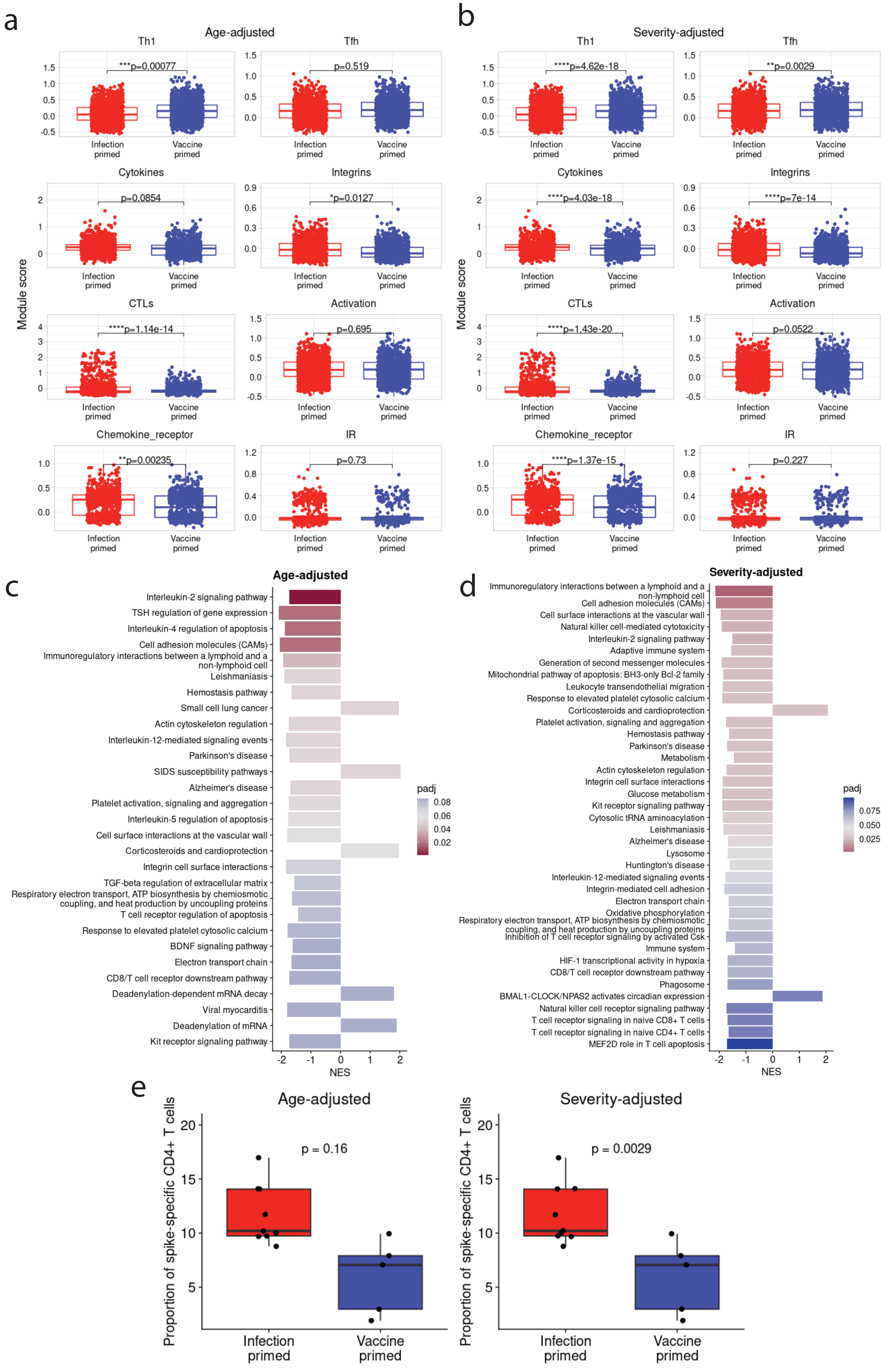

### Supplemental Figure 5

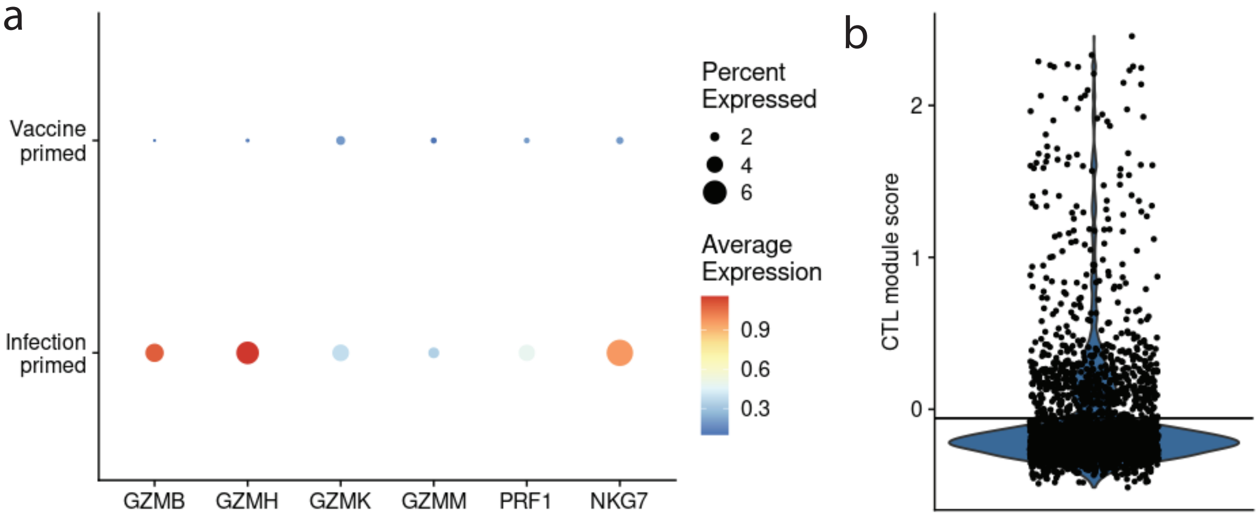
